# A high-resolution, unbiased analysis of the cellular immune response to Epstein-Barr virus

**DOI:** 10.1101/2025.10.09.681317

**Authors:** Álvaro F. García-Jiménez, Arianna Picozzi, Andrea Sánchez de la Cruz, Enrique Vázquez, Alberto Benguría, Ana Dopazo, Vassilios Lougaris, Luis Ignacio González-Granado, Obinna Chijioke, Eduardo López-Granados, Hugh T. Reyburn

## Abstract

More than 95% of humans are infected with Epstein-Barr virus (EBV), yet although EBV infection has been associated with inflammatory and autoimmune diseases, lymphoproliferative disorders, and several types of cancer, for the vast majority of infected people the infection is asymptomatic as EBV replication is controlled by the immune system. Immunity against this virus has been studied since the discovery of EBV in the 1960s, and although important insights have been made, no unbiased, global studies of immune responses to EBV in healthy seropositive subjects have been reported. Here we describe a novel protocol to study the cellular immune response to EBV, detecting lymphocytes that respond to EBV via analyses of proliferation or induced expression of activation markers and cytokines. Using this system we sequenced, for the first time at a single-cell level, the transcriptome of all cells capable of responding to EBV in healthy individuals and in patients with inborn errors of immunity (IEI) associated with susceptibility to EBV infection. Lymphocyte cytotoxicity appears to be crucial for the proper control of EBV-infection, while a proportionate T-regulatory cell response likely helps to avoid excessive immunity and immune pathology. We also show that γδ T cells expressing the TCR Vδ1 chain use various activating natural killer (NK) cell receptors to recognise and kill EBV-infected lymphoblastoid cell lines (LCLs) and thus could be a promising candidate for allogeneic cell therapy for EBV-associated lymphoproliferative disorders in patients with either primary or secondary immunodeficiencies.

**Graphical abstract:** 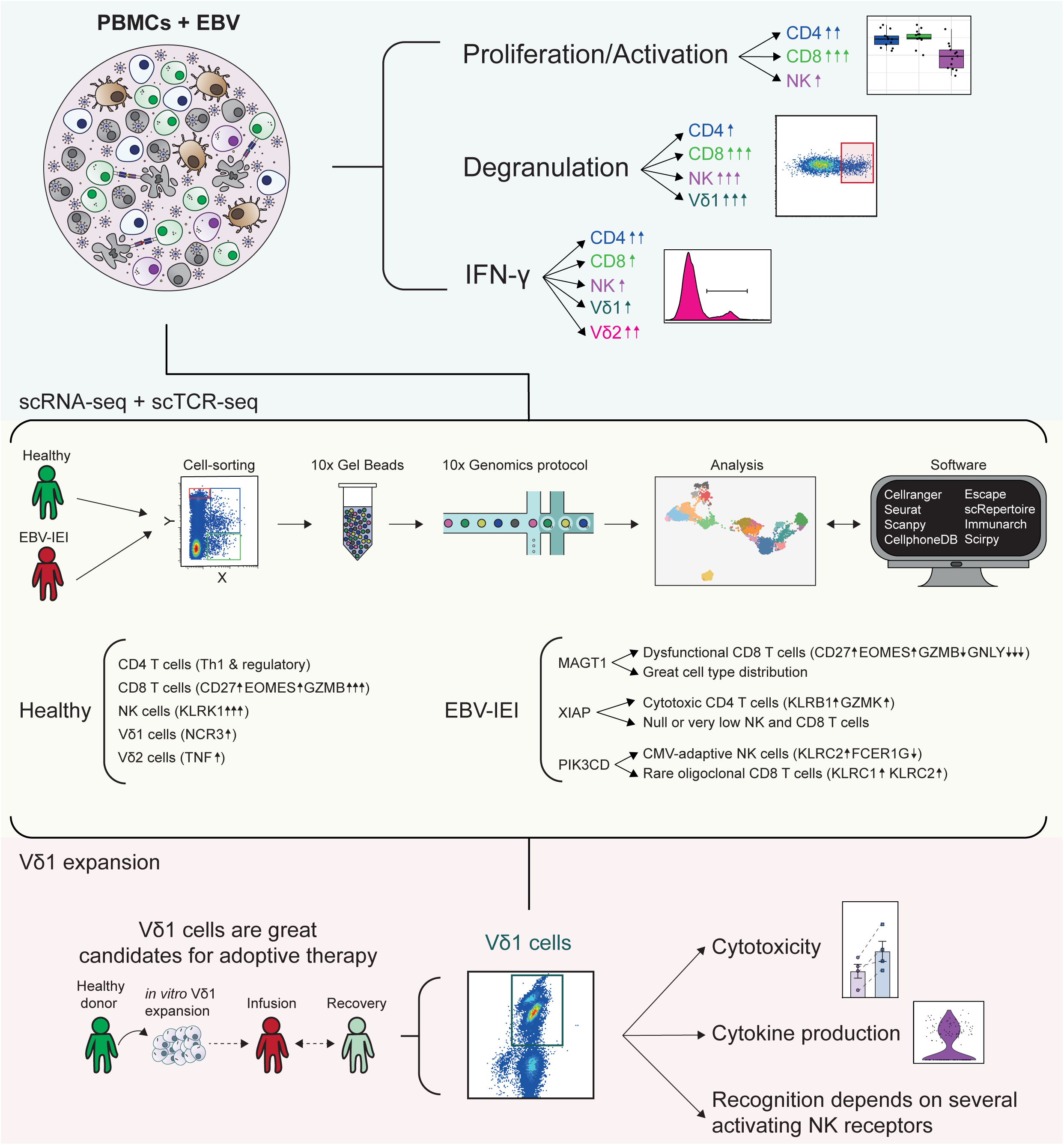

## INTRODUCTION

The Epstein-Barr virus (EBV) is a ubiquitous herpesvirus that is primarily transmitted through saliva and infects more than 95% of the adult human population worldwide^1^. Indeed, EBV has coevolved with humans and the result of this process is a finely balanced relationship where in healthy individuals first infected as children, the initial exposure to EBV generally does not lead to significant complications, and the majority of the population experiences an asymptomatic infection^1,2^. However, if the infection occurs during adolescence, approximately 15% of people experience a mild febrile illness known as infectious mononucleosis (IM)^2,3^. Given the seroprevalence of the virus, infectious mononucleosis occurs relatively frequently and, over the years, much of the immunological research on EBV has focussed on this model of primary infection even though IM is actually a pathological situation where an exaggerated response of CD8 T cells underlies the development of the disease^3^. EBV infection can also lead to the development of other diseases. EBV was the first human oncovirus to be identified and is etiologically linked to around ∼2% of all human cancers (some 300,000 new cases and 200,000 cancer deaths every year)^4^. These include several B cell malignancies: Burkitt lymphoma, classic Hodgkin lymphoma, and post-transplant lymphoproliferative disorders; extranodal lymphomas of T or natural killer (T/NK) cell origin, as well as undifferentiated nasopharyngeal carcinoma; smooth muscle cell sarcomas, and some gastric carcinomas^4,5^. Further, compelling epidemiological data also implicate EBV infection as a key trigger for the development of multiple sclerosis (MS)^6^, the most common cause of nontraumatic neurological disability in young adults^7^. Finally, EBV is also a major cause of morbidity and mortality in individuals with a range of inborn errors of immunity (IEI). Mutations in multiple genes, including *MAGT1*, *XIAP*, *SH2D1A*, *CD27*, *CD70*, *ITK*, *RASGRP1*, *CTPS1*, *CORO1A*, *TNFRSF9*, *PRF1*, *RAB27A*, *DEF6*, *STX11*, *PIK3R1*, *UNC13D*, *STXBP2*, *ORAI1*, *DOCK8*, *FCGR3A*, *LYST*, *TNFSF9*, or *PIK3CD* have been associated with enhanced susceptibility to EBV disease^8,9^. These studies have also provided important insights into the immune response to EBV, confirming the importance of EBV-specific CD8 T cells and NK cells for immune control of EBV-infected B cells. However, since specific inborn errors of immunity are extremely rare diseases, the characterisation of immune populations in these patients necessarily involves study of only a small number of patients. Moreover, in most cases, multiple immune cell lineages express the affected genes and this makes it difficult to relate how the mutation confers susceptibility to EBV disease with changes in specific populations.

The immune response made by persistently infected, but healthy seropositive individuals, the situation in which host immunosurveillance efficiently controls viral replication, has been studied less, but is known to include innate immunity, EBV-specific CD8^+^ T-cell responses to lytic and latent cycle antigens and a lesser, often 10- fold lower, EBV-specific CD4^+^ T-cell response as well as the production of virus-neutralising antibodies^3,10^.

Data from studies of infectious mononucleosis show that the activation and proliferation of natural killer (NK) cells seem to aid control of EBV infection since NK cell numbers are significantly elevated at diagnosis of IM and these higher NK cell counts associate with significantly lower viral loads in peripheral blood^11^. Moreover, it has been demonstrated that the CD56^bright^ subset prevents B-cell transformation via the secretion of IFN-γ, and that NK cells of an early differentiated phenotype (CD56^dim^NKG2A^+^KIR^-^) act rapidly to restrain the lytic cycle^12,13^. Interestingly, *in vitro* studies using models of LCL stimulation found that those individuals who did not make a marked NK cell proliferative response instead expanded Vγ9Vδ2 gamma-delta T cells^14^. Although the cytotoxic capacity of these cells is controversial, Vγ9Vδ2 T cells can produce various cytokines, including IFN-γ or TNF-α, that might help to promote an effective immune response against EBV^15^.

The adaptive immune response plays a crucial role in controlling viral replication and preventing EBV-associated diseases. CD8 T cells are particularly important in eliminating infected cells, as numerous studies have shown significant clonal expansion of CD8 T cells during infectious mononucleosis, and a rapid recall after re-exposure to EBV^16^. This response is more pronounced in the tonsils than in the blood, although the TCR repertoire is similar in both locations, indicating non-selective recruitment. Moreover, it is estimated that only 10% of the unique clonotypes detected during infectious mononucleosis are detected again during convalescence, suggesting continuous renewal of the CD8 T cell immune repertoire during persistent infection^17^. These cells are differentiated effectors expressing T-bet or Eomes, and they are believed to play a crucial role in controlling EBV reactivation, as many of the IEI-causing mutations associated with EBV-related lymphoproliferative disorders lead to problems with the proper function of CD8 lymphocytes^18^. On the other hand, the CD4 T cells detected in experiments of infection with EBV include central memory CD4 T cells (CCR7^+^) or effector memory CD4 T cells (CCR7^-^) with a predominant Th1 phenotype and increased secretion of TNF-alpha^19^. In persistent infection, regulatory CD4 T cells (FOXP3^+^) also gain importance and are maintained in the tonsils at levels similar to those observed during infectious mononucleosis^20^. Although the CD4-type response has always been in the background of research on the immune response to EBV, it seems to play a crucial role in immune regulation, targeting latently infected cells, or providing B cell help. In fact, the humoral response is of particular importance in the immunosurveillance of infected cells and recent studies suggest that antibodies against lytic antigens, such as BFRF3, BLRF2, and BdRF1, are actually dominant in the antibody response to EBV^21^. However, currently, a global view of how these different aspects of immunity to EBV are integrated is lacking. To achieve a better understanding of the spectrum of cellular components involved in the control of EBV infection, we have used scRNA-seq and scTCR-seq to characterise the transcriptome and clonotypes of immune cells from healthy asymptomatic subjects responding to EBV, and complemented these data in flow cytometry and functional assays. In parallel, we have also analysed the immune responses made by patients with genetic defects known to confer susceptibility to EBV-mediated pathology. Our hypothesis is that the joint study of patients with genetic defects that lead to inefficient control of EBV replication *in vivo*, coupled with characterisation of the immune response in healthy individuals, could lead to a better definition of the molecular and cellular aspects of the immune response by which an individual can cope with persistent EBV infection. Indeed, despite all the work carried out throughout the years, few studies have been dedicated to high-resolution characterisation of the major memory immune populations that react against EBV. During the course of these experiments we also discovered that γδ T cells expressing the TCR Vδ1 chain use various activating NK cell receptors, not the TCR, to recognise and kill EBV transformed B cells. In experimental animal models we have gone on to show that TCR Vδ1 T cells, when transferred to mice burdened with an EBV LCL tumour, preferentially localise to the tumour and show some promise as an allogeneic therapy for EBV-associated lymphoid tumours.

## RESULTS

### 1. Development of an in vitro system to study the EBV-specific cellular immune response in peripheral blood

To establish a minimally biased model to study cellular immunity to EBV, we isolated PBMCs from peripheral blood since blood contains a great variety of effector immune cells, as well as B cells, that are physiologically relevant targets for EBV infection. The B95-8 strain of EBV was used to infect PBMCs *in vitro*. This treatment is known to activate latently infected B cells to move into productive replication as well as infecting previously uninfected-cells so that infected B cells in the culture express a full range of lytic and latent-phase viral antigens derived from both endogenous and exogenous EBV^22^.

To test whether the immune cells activated by this protocol were enriched for EBV- reactivity, restimulation experiments were carried out. On day 0 of these assays, donor PBMC reactivity to pools of peptide epitopes from HCMV and EBV was evaluated in short-term assays of IFN-γ and TNF-α production. In parallel, another aliquot of PBMCs were co-cultured with EBV-infected autologous PBMCs. After 7 days, these cultures were supplemented with IL-2 and allowed to rest for 5 more days before being challenged with pools of either HCMV or EBV peptide epitopes. Comparison of the day 0 and day 12 data demonstrate that T cells producing IFN-γ and TNF-α after restimulation with EBV, but not HCMV, peptide pools had been specifically enriched by prior exposure to EBV-infected PBMCs, proving that this method preferentially expanded EBV-specific clones (**Figure 1A**). When PBMCs from XMEN and XIAP_1 patients were tested in these assays, initial stimulation with EBV followed by restimulation with EBV peptides led to marked IFN-γ accumulation by CD8 T cells from XMEN, while it was modest and focussed on CD4 T cells in the case of XIAP_1 (**Supplementary Figure 1A,B**).

**Figure 1.**
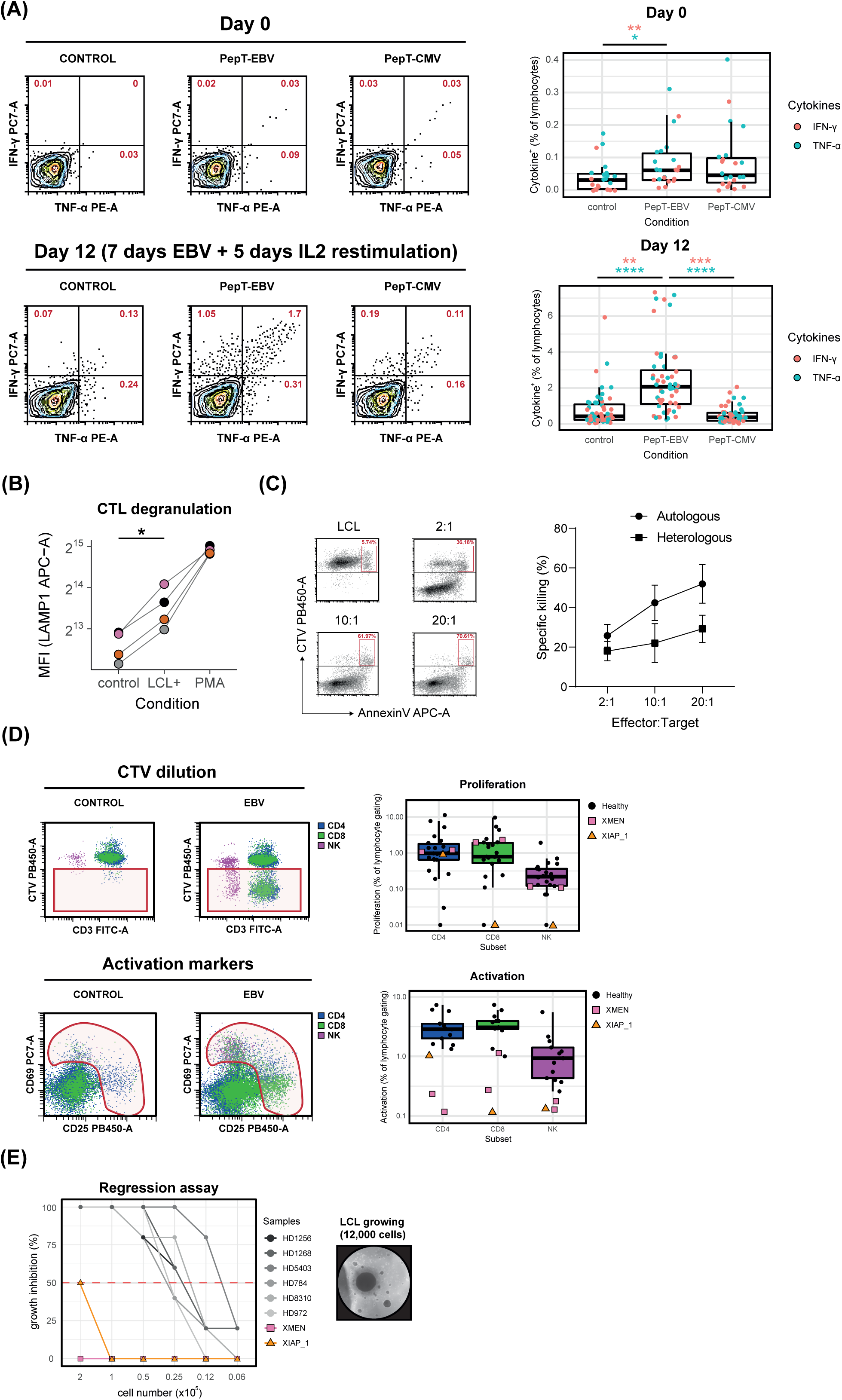
*In vitro* detection of immune cell reactivity against EBV in peripheral blood. (A) Identification of IFN-γ and TNF-α producing T cells from healthy samples after incubation with EBV or HCMV peptides at day 0 or after incubation with EBV for 7 days, rest with IL-2 for 5 days, and restimulate with EBV or HCMV peptides at day 12. One-way ANOVA with Tukey post-hoc tests (*p<0.05, **p<0.01, ***p<0.001, ****p<0.0001). (B) Upregulation of LAMP1 in healthy donor CTLs after incubation with autologous LCLs or PMA/ionomycin. Paired sample t-test (*p<0.05). (C) Specific-killing assays between healthy donor CTLs and autologous or heterologous LCLs at different effector:target ratios. (D) Assessment of proliferation or activation of CD4, CD8, or NK cells after EBV incubation for 6 days in healthy and IEI samples. (E) Regression assays representing the growth inhibition rate of monocyte-depleted PBMCs (6 healthy donor, XMEN and XIAP_1 samples) after EBV incubation during 4 weeks at different confluence levels. A photograph of growing LCLs is included.

To further characterise EBV-specific immunity, lines of cytotoxic T lymphocytes (CTLs) were prepared from peripheral blood lymphocytes by cycles of stimulation with autologous lymphoblastoid cell lines (LCLs) followed by IL-2 supplementation. Cytotoxic lymphocytes generated from healthy donors, mainly consisted of CD8 T cells that upregulated the degranulation marker LAMP1 upon exposure to autologous LCLs and efficiently killed autologous or heterologous LCLs over a range of effector:target ratios (**Figure 1B,C, Supplementary Figure 1C**). Similar findings were described in XMEN patients, in which the CTLs, mainly CD8^+^, could upregulate LAMP1 after stimulation with autologous LCLs or PMA/ionomycin, and also showed some low degree of specific cell killing of autologous LCLs (**Supplementary Figure 1D-F**). In contrast, CTLs could not be generated from PBMCs of the XIAP_1 patient.

Finally, EBV-reactive immune cells were identified either by the induced expression of activation markers or by the dilution of CTV-fluorescence in proliferating cells. Quantitative analysis for responses of CD4 and CD8 T cells as well as NK cells from healthy and immunodeficient donors are shown in **Figure 1D**. These experiments suggest, in contrast to previous studies^3,10^, that the absolute frequencies of responding CD4 and CD8 T cells seem to be comparable in healthy samples. Moreover, XMEN and XIAP_1 patients were capable of mounting a detectable immune response, but it was generally weaker than those made by healthy donors.

Strikingly, however, although PBMCs from XMEN and XIAP_1 patients could recognise EBV infected cells, PBMCs from these patients were unable to control B cell transformation by EBV. LCL outgrowth was efficiently controlled in PBMC cultures from healthy donors even at low cell densities, whereas PBMCs from XMEN or XIAP patients either only reduced, or completely failed to limit, the establishment of EBV-transformed LCLs (**Figure 1E**).

### 2. scRNA-seq shows the proliferation of infected B cells and identifies EBV-responsive immune cell populations in healthy and immunodeficient individuals

In order to analyse in detail qualitative aspects of the cellular immune response to EBV infection made by PBMCs from healthy compared to immunodeficient donors, the single-cell transcriptome and TCR usage of PBMCs proliferating after exposure to EBV was characterised (**Supplementary Figure 2A**). 4 samples from healthy subjects were studied, including 1 sample that responded against HCMV but not EBV, that was termed “EBV-naive” (**Supplementary Figure 2B**). 4 samples from immunodeficient patients [3 MAGT1^y/-^ (XMEN_1, XMEN_2, XMEN_3) and 1 XIAP^y/-^ (XIAP_1)] were analysed in parallel. Additionally, naive/resting cells from the negative fraction of healthy samples were also loaded to establish a basal transcriptome for non-responding cells.

We sequenced 7,250 cells that were grouped in 18 clusters and 6 general identities based on the differential expression of gene markers (**Figure 2A,B**). These clusters included B and plasma cells, monocytes, CD4 T cells, cytotoxic cells (comprising CD8αβ and γδ T cells as well as NK cells), and the naive/resting cells of the negative fraction. Strikingly, an increased proportion of CD4 and cytotoxic cells were found in samples from healthy donors, whereas samples from immunodeficient and the EBV- naive donors were enriched for clusters of the B cell compartment (**Figure 2C, Supplementary Figure 2C, Supplementary Table 1**). However, this analysis did identify some expansion of effector cells in samples from the immunodeficient patients that were not detected in the EBV-naive control (e.g., C3, C7, C9), again supporting the specificity of the working model and the ability of MAGT1^y/-^ and XIAP^y/-^ lymphocytes to respond to EBV infection.

**Figure 2.**
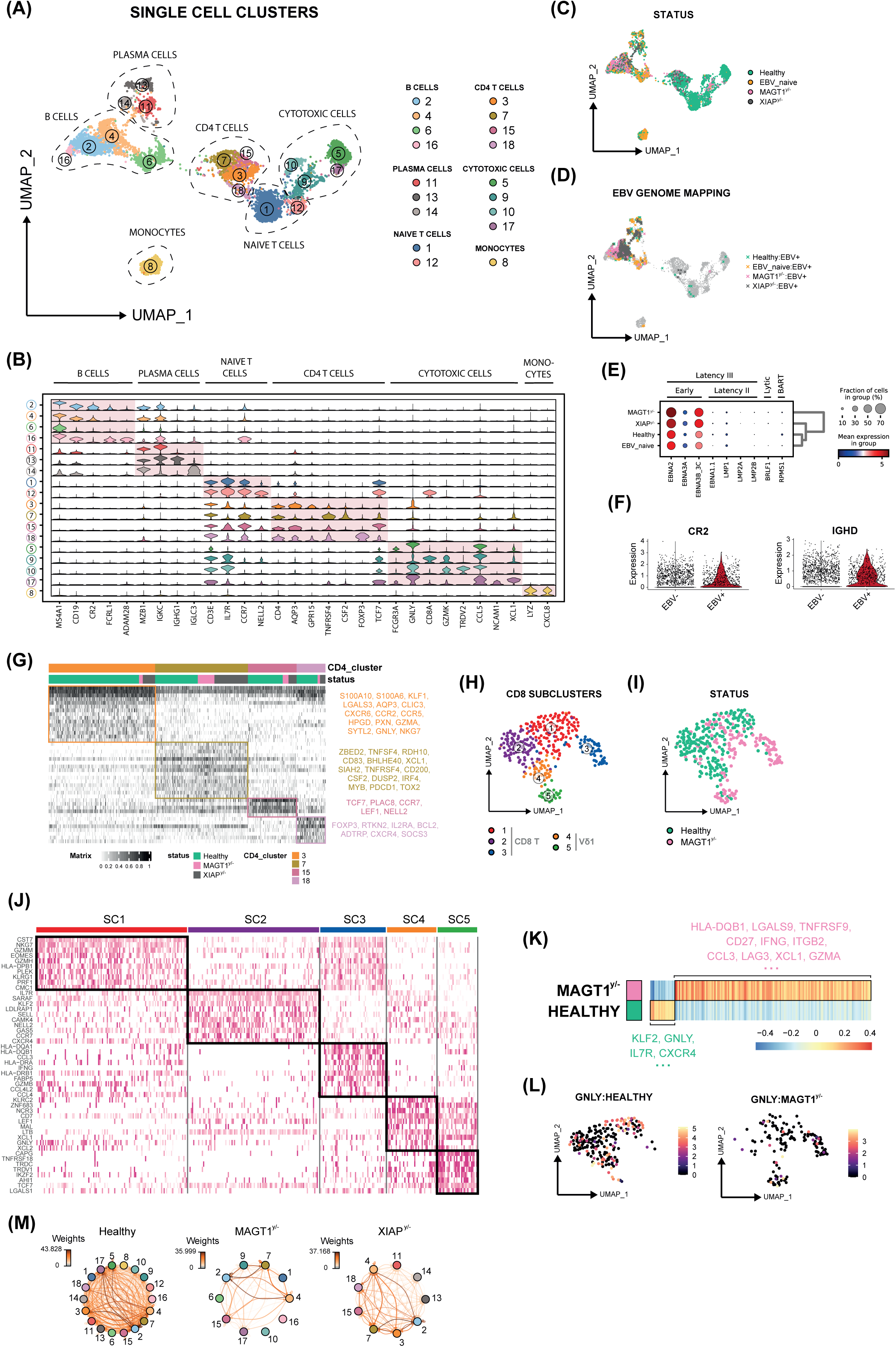
Transcriptomic analysis of cells proliferating after exposure to EBV. (A) UMAP representation of the whole dataset clustered at 1.2 resolution and grouped in general identities. (B) Stacked violin-plot showing differential gene markers of each cluster. (C) UMAP distribution per status. (D) UMAP showing cell points with detection of EBV transcripts per status. (E) Dot-plot of the EBV detected genes. (F) Violin-plot showing the expression of *CR2* and *IGHD* genes within infected and non-infected cells. (G) Scaled heatmap of the most differential expressed genes within CD4 clusters. (H) UMAP representation of C9 subclustered at 0.5 resolution. (I) C9 UMAP distribution per status. (J) Heatmap showing the top10 differentially expressed genes per C9 subcluster. (K) C9 pseudobulk heatmap representing the scaled mean expression values of differentially expressed genes per status. (L) Feature plots coloured by GNLY expression for healthy and MAGT1^y/-^ cells within C9. (M) Differential cell-cell interactions between clusters per status. Clusters with < 20 cells were omitted.

When reads were aligned against the EBV genome, EBV transcripts were mainly detected in B (C2, C4) and plasma cell (C11, C13, C14) compartments (**Figure 2D**), except for MAGT1^y/-^ individuals, where plasma cells were not detected. Moreover, the numbers of positive events were ∼3 times lower in healthy samples if compared with samples from the EBV-naive or immunodeficient subjects (**Supplementary Figure 2D**). In general, EBV positive cells expressed the viral *EBNA2*, *EBNA3A*, and *EBNA3B/C* genes, indicative of the first steps of EBV transformation (**Figure 2E**), and also expressed the human genes *CR2* and *IGHD*, that are the viral receptor and a marker of naive B cell identity (**Figure 2F**).

CD4 T cells were divided in 4 clusters (C3, C7, C15, C18) and were captured in MAGT1^y/-^ and XIAP^y/-^ samples (**Figure 2G, Supplementary Figure 3A**). C15 and C18 represented subpopulations with a resting (*TCF7*, *CCR7*) and a regulatory (*FOXP3*, *IL2RA*) phenotype, respectively. C3 was predominantly found in healthy samples, and comprised a mature cluster with expression of homing (*CXCR6*, *CCR2*) and granulocytic (*GNLY*, *SYTL2*) markers. In contrast, C7 appeared enriched in the samples from IEI patients, and was defined by the transcription factor *ZBED2*, with evidence of *TNFSF4*/*TNFRSF4* co-expression. These cells also expressed *TOX2*, that has been shown to increase central memory CAR T cell differentiation, but not proliferation, because of upregulation of exhaustion precursor pathways^23^. Clonal expansion of TCRαβ expressing cells was a striking feature of the C3 cluster, in healthy seropositive donors, but this was less prominent in the patients. (**Supplementary Figure 3B**,C).

Cytotoxic cells also grouped in 4 major clusters (C5, C9, C10, C17) that were detected in healthy and MAGT1^y/-^ individuals, but not in XIAP_1 (**Figure 2C**). C9 was enriched in CD8^+^ T cells and could be reclustered in 5 well-defined identities despite the low number of cells (**Figure 2H,I**). Subclusters 1 and 3 were highly cytotoxic cells (*KLRG1*, *GZMM*, *EOMES*) that probably represent a mix of mature and immature cells since *FCGR3A* is transiently detected (**Supplementary Figure 3D**), SC2 comprised resting mature cells (*IL7R*, *KLF2*) that were not detected in MAGT1^y/-^, and SC4-5 expressed markers of Vδ1 identity (*TRDV1*, *IKZF2*, *KLRC2*) (**Figure 2J**). Pseudobulk analysis of C9 demonstrated the upregulation of activation markers in MAGT1^y/-^ (*TNFRSF9*, *CD27*), but lower expression of *GNLY* or *CXCR4* compared with healthy samples (**Figure 2K,L**). Clonal expansion of C9 subclusters was inexistent in SC4-5, and limited in SC1 and SC2, probably as a consequence of low cell detection (**Supplementary Figure 3E**,F). Interestingly, there were great differences in the distribution of subclusters between healthy and MAGT1^y/-^ samples (**Figure 2I, Supplementary Figure 3G**), but these differences might be due to different states of activation/proliferation between subclusters 1-3 and 4-5, as depicted by Pearson’s correlation and expression of *MKI67* (**Supplementary Figure 3H**,I).

The C5 and C17 clusters were identified as CD56^dim^ (*CX3CR1*, *FGFBP2*, *FCGR3A*) and CD56^bright^ (*NCAM1*, *COTL1*, *XCL1*) NK cells, whereas C10 comprised a mixture of MAIT (*TRAV1-2*, *KLRB1*) and Vδ2 (*TRDV2*, *TRGV9*) T cells (**Supplementary Figure 3J**,K).

Overall, the transcriptomic profiles of proliferating cells identified cell clusters (C3, C5, C7, C9, C17, C18) with great potential to directly interact with EBV-infected B cells (C2, C4) in healthy, MAGT1^y/-^, and XIAP^y/-^ samples (**Figure 2M**).

### 3. Single-cell characterisation of the transcriptome and TCR of activated immune cells highlights important aspects of cellular immunity to EBV

The previous unbiased scRNA-seq experiment helped to draw a general landscape of the cellular immune response to EBV. However, to capture similar cell states between conditions and better explore intra-specific differences between clusters, we repeated these experiments analysing the single-cell transcriptome and TCRs, selecting immune cells responding to EBV on the basis of induced expression of the activation markers CD69 and/or CD25 after exposure to EBV (**Supplementary Figure 4A**). Additionally, in this second set of experiments, B cells were depleted from patient samples, via a CD19 negative selection step, to increase the representation of cells responding to EBV. The samples analysed included 2 of the previous related MAGT1^y/-^ donors (XMEN_1, XMEN_3), 1 new XIAP^y/-^ donor (XIAP_2), and 2 unrelated PIK3CD^GOF^ donors (PIK3CD_1, PIK3CD_2), respectively. Samples from 9 new healthy donors were also analysed to gain more insight into the immune cell subpopulations that emerged during the response to EBV infection. EBV-reactive cells were sorted from PBMCs at day 6 since the activation markers were still detected in healthy samples and very little upregulation of the exhaustion marker PD-1 was noted in CD8 T cells (**Supplementary Figure 4B**,C).

A total of 39,559 cells from the different donors were integrated *in silico* to obtain a UMAP landscape of 21 clusters, that allowed the identification of some general populations like CD4 T cells, CD8 T cells, NK cells, MAIT and Vδ2 cells, or B and plasma cells (the latter only present in healthy samples) (**Figure 3A, Supplementary Figure 4D**). The distribution of cells per condition was clearly heterogeneous. Samples from healthy and MAGT1^y/-^ donors occupied essentially the whole 2D UMAP space, whereas the XIAP^y/-^ sample was enriched in CD4 T cells (78.3%), and the PIK3CD^GOF^ samples contained a majority of either CD8 T cells (49.3%:PIK3CD_1) or NK cells (38.3%:PIK3CD_2) (**Figure 3B, Supplementary Table 2**)., A pseudobulk principal component analysis (PCA) was performed on the dataset (excluding plasma and B cells), to visualise the general differences within the whole detected gene expression values between the samples. Immunodeficient samples were clearly divided in two groups (XMEN_1:XMEN_3 and XIAP_2:PIK3CD_1:PIK3CD_2), whereas healthy samples were separated from deficient samples but with clear differences in intragroup colocalisation, probably as a consequence of the heterogeneity in the magnitude of the response to EBV and as a consequence of batching (**Supplementary Figure 4E**).

**Figure 3.**
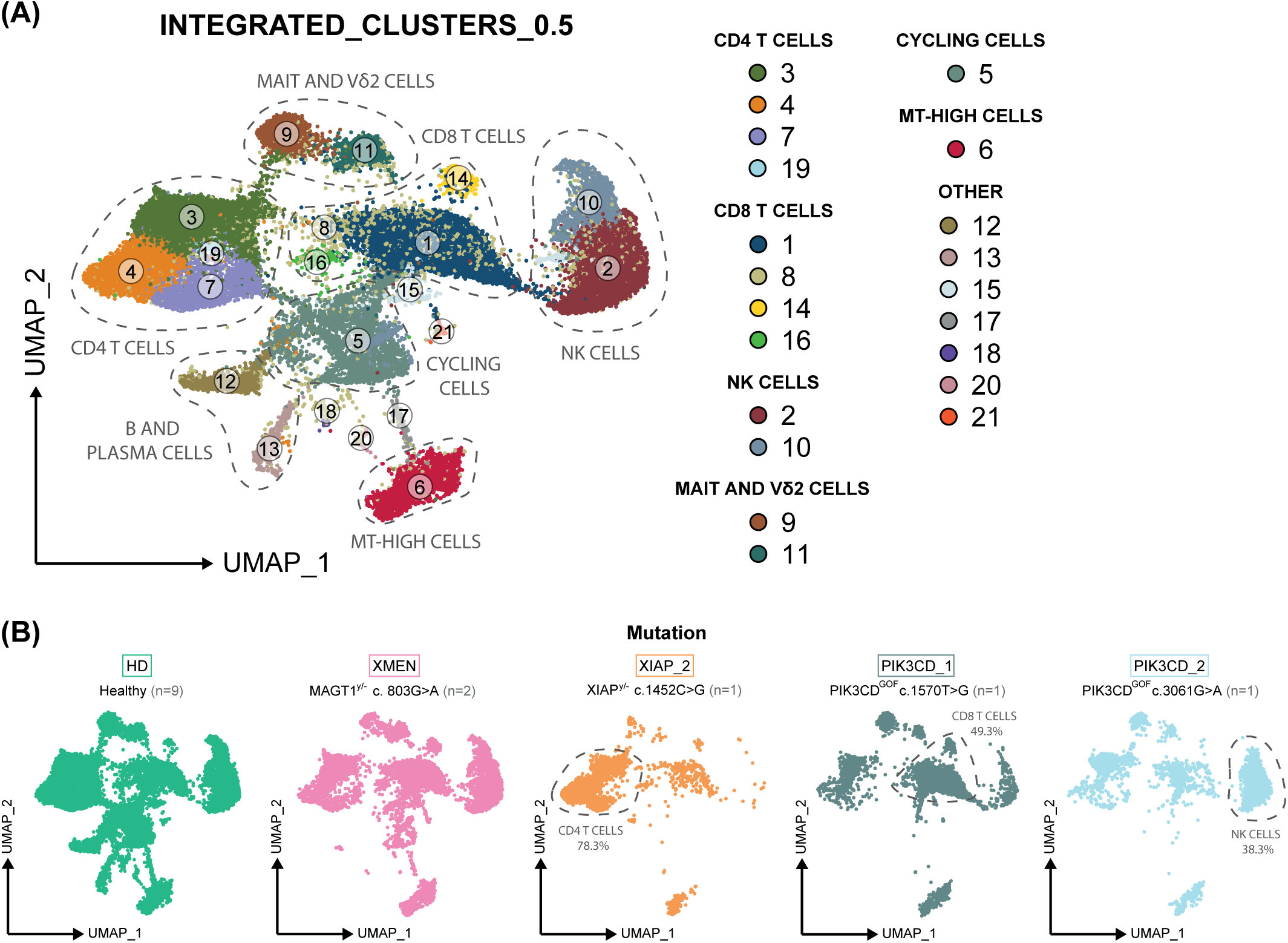
Clusterisation of cells activated by exposure to EBV. (A) UMAP distribution of the integrated cells from different experiments at 0.5 resolution. (B) UMAP split and coloured by mutation.

#### 3.1. CD4 T cells

Non-naive CD4 T cells were divided in 7 subclusters that were differentially distributed between cells from healthy subjects and XMEN, XIAP_2, and PIK3CD^GOF^ patients (**Figure 4A,B, Supplementary Figure 5A**). Regulatory T cells (*IL2RA*^+^*FOXP3*^+^) were markedly more abundant in healthy donors compared to patients. Moreover, two distinct subclusters of regulatory T cells (SC4, SC6) could be clearly distinguished. One of them (SC6) was enriched in molecules characteristic of activated cells (*HLA-DRB1*, *CCR4*, *TNFRSF18*) (**Figure 4C**), whereas the subcluster 4 has more of a resting phenotype and its presence may be a consequence of the use of CD25 in the sorting strategy. Strikingly, the major populations of effector CD4 T cells also varied notably between healthy donors and IEI patients. SC5 cells expressing chemokine receptors like *CXCR3* or *CCR2* together with cytotoxic and effector molecules such as *GZMB*, *GNLY*, or *CSF2* were abundant in healthy donors. The SC3 cells, present in XMEN patients, displayed a resting phenotype with marked expression of *IL7R*, type I IFN-induced molecules, *TNFSF10*, and little cytotoxic potential, whereas the SC2 cells, mainly found in XIAP deficiency, were somewhat similar to healthy cells, showing a more cytotoxic phenotype with expression of *KLRB1* and *GZMK* (**Figure 4D, Supplementary Figure 5B**). Interestingly, given the inflammatory phenotype of XIAP- deficiency, two recent studies have shown that T cells expressing GZMK are found in inflamed tissues, where they contribute to disease pathology by driving complement activation in a tissue-restricted way^24,25^. Multiple expanded clones of CD4 T cells were present in SC5, but generally the extent of these expansions was moderate (<10), suggesting a polyclonal CD4 response in healthy samples. A few expanded clones were also observed in IEI samples (**Figure 4E, Supplementary Figure 5C**). Overall, these data suggest that CD4 T cells polarised into Th1 with cytotoxic potential and CD4 T_reg_ cells are two key components of a balanced CD4 T cell response to EBV infection in healthy subjects.

**Figure 4.**
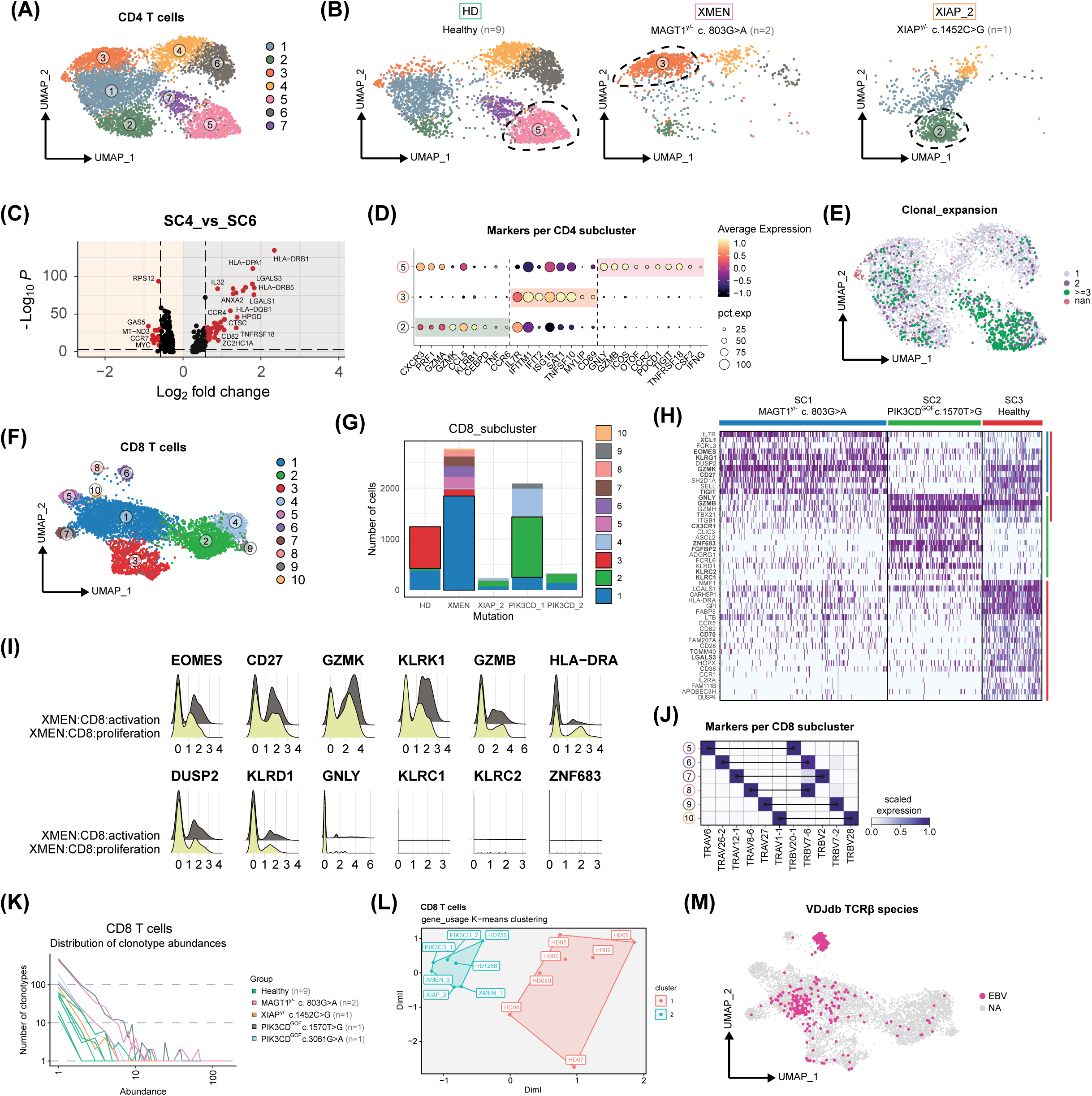
Transcriptomic analysis of adaptive cells activated against EBV. (A) UMAP distribution of CD4 T cells at 0.5 resolution. (B) UMAP distribution of CD4 T cells split by mutation. (C) Volcano plot representing the genes differentially expressed between SC4 and SC6 within CD4 T cells, highlighting in red genes with a Log2FC > 0.58 and a Bonferroni adjusted p-value < 10^−4^. (D) Dot plot of different markers that define SC2, SC3, and SC5 in CD4 T cells. The scaled average expression is represented in a colour density, and the percentage of cells expressing a specific marker is represented through the dot diameter. (E) UMAP coloured by clonal expansion within CD4 T cells. (F) UMAP distribution of CD8 T cells at 0.5 resolution. (G) Stacked bar-plot showing the distribution of CD8 T cell subclusters per mutation. (H) Heatmap showing differentially expressed genes between SC1, SC2, and SC3 within CD8 T cells. (I) Ridge-plots of some markers present or absent in the CD8 T cell fractions of XMEN samples from proliferation and activation scRNA-seq experiments. (J) Matrix plot of *TRAV* and *TRBV* gene detection in CD8 T cell SC5-SC10. (K) Distribution of clonotype abundances per mutation within CD8 T cells. (L) K-mean clustering of samples according to the VDJ usage. (M) UMAP coloured by EBV TCRβ specificity according to VDJdb.

#### 3.2. CD8 T cells

When clustering analysis focussing on CD8 T cells was performed, 10 subclusters were identified in cells from healthy, XMEN, and PIK3CD_1 samples (**Figure 4F,G**). Most of the cells captured in XMEN:SC1 and Healthy:SC3 were *KLRG1*^+^*CD27*^+^*TIGIT*^+^, with consistent expression of *GZMK*, but low expression of *GZMB*, *GZMH*, and especially *GNLY,* in the case of the XMEN subjects, similar to what was seen in the proliferation-based scRNA-seq (**Figure 4H,I**). The SC5-SC10 clusters in the UMAP of CD8 T cells were essentially defined by TRAV and TRBV chain usage, representing examples of highly expanded clones that were mainly detected in MAGT1^y/-^ patients (**Figure 4J, Supplementary Figure 5D**). Curiously, some of these clonal subclusters corresponded to *KLRC2*^+^*GZMB*^+^*GNLY*^+^ cells that upregulated innate molecules like *XCL1*, *KLRF1*, or *KLRC3* (**Supplementary Figure 5E**). Globally, the analysis of CDR3 sequences revealed that individual clones could be detected more than ten times in many donors, and the analysis of VJ and VDJ usage revealed a marked TCRαβ restriction in immunodeficient samples compared to healthy samples, as also demonstrated when analysed using the “Grouping of Lymphocyte Interactions by Paratope Hotspots” (GLIPH) algorithm (**Figure 4K,L, Supplementary Figure 5F**,G, **Table 1**). Interestingly, we found several examples of expanded clones in our dataset where the TCR had already been annotated as EBV-specific in the VDJdb database, particularly in the MAGT1^y/-^ samples (**Figure 4M, Supplementary Figure 5H**). Very few CD8 expressing lymphocytes were detected in samples from the XIAP_2 or the PIK3CD_2 patients (**Figure 4G**), while the majority of CD8 expressing cells detected in PIK3CD_1:SC2 were *ZNF683*^+^*FGFBP2*^+^*CD27*^-^ with marked expression of killer cell lectin-like receptors (*KLRD1*, *KLRC1*, *KLRC2*), marking some kind of resident phenotype in peripheral blood (**Figure 4H**).

**Table 1.**
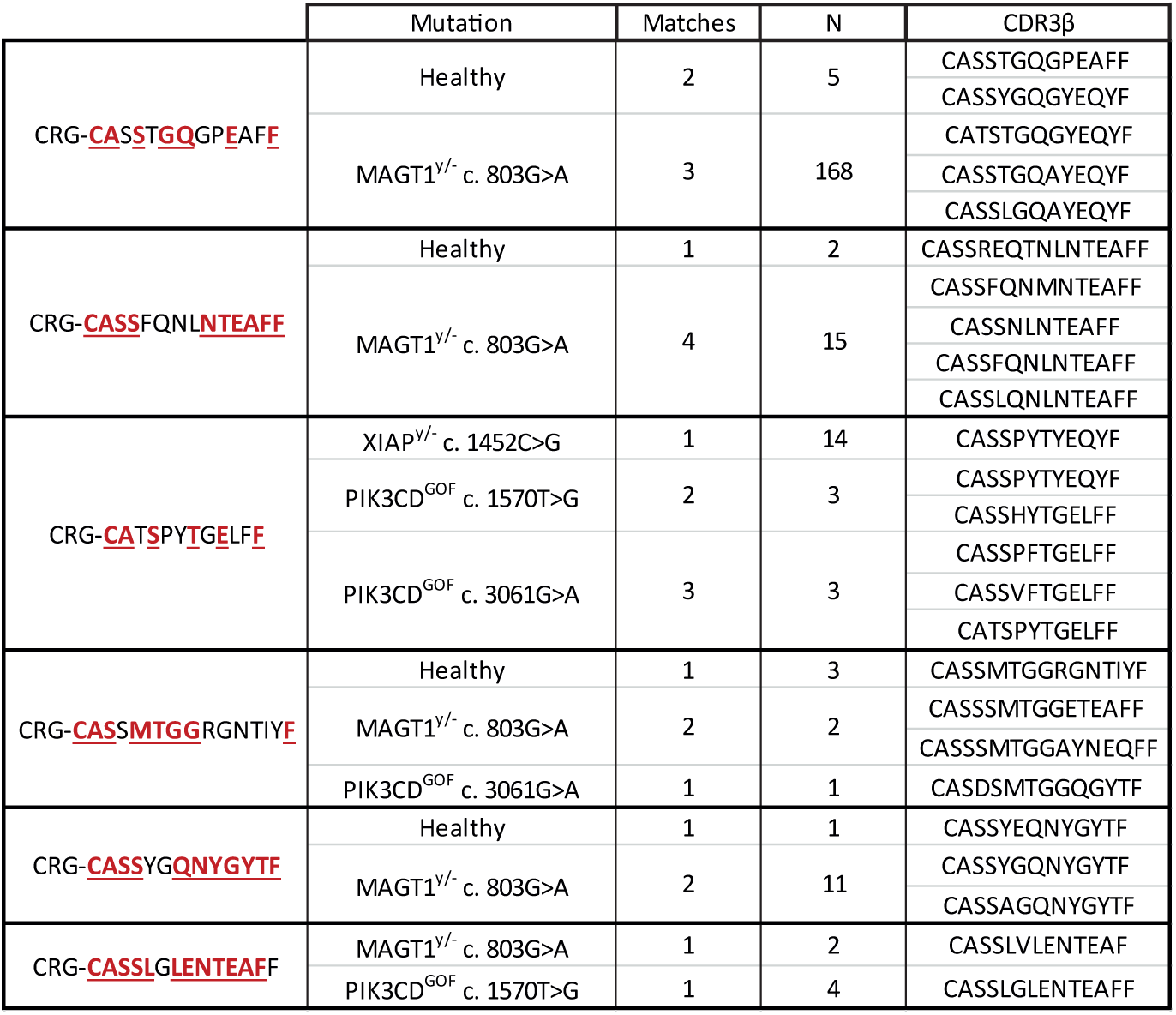
Example of CD8αβ convergence groups identified with the GLIPH algorithm. CDR3βs found to be similar were grouped and stratified per mutation. The column “Matches” represented the number of different CDR3βs identified within a convergence group, and the column “N” the total number of cells that carried these CDR3βs.

#### 3.3. Innate lymphocytes

Most NK cells present in healthy and XMEN samples were of the CD56^dim^ phenotype (**Figure 5A, Supplementary Figure 6A**-C). These CD56^dim^ NK cells did not express the majority of *KIR* genes, but were positive for *FCGR3A*, *KLRC1*, or *KLRK1*, although MAGT1^y/-^ patients displayed a somewhat more effector phenotype (*CX3CR1*, *CCL4*, *GZMB*, *GZMH*) with decreased expression of *CXCR4*, *JUN*, *LTB*, or *XCL2* (**Figure 5B, Supplementary Figure 6D**). The proportions and phenotypes of CD56^bright^ NK cells were quite similar between healthy and XMEN conditions, but the latter appeared more active and proliferative, given the expression of *CD38* and *STMN1*, but with lower expression of *IL7R*, and undetectable *JAML* (**Figure 5C**). Again, very few NK cells were found in the XIAP_2 and PIK3CD_1 patients but, curiously, an expansion of HCMV-associated adaptive NK cells (*KLRC2*^+^*FCER1G*^-^*KLRC1*^-^) was observed in the PIK3CD_2 donor (**Figure 5A, Supplementary Figure 6E**).

**Figure 5.**
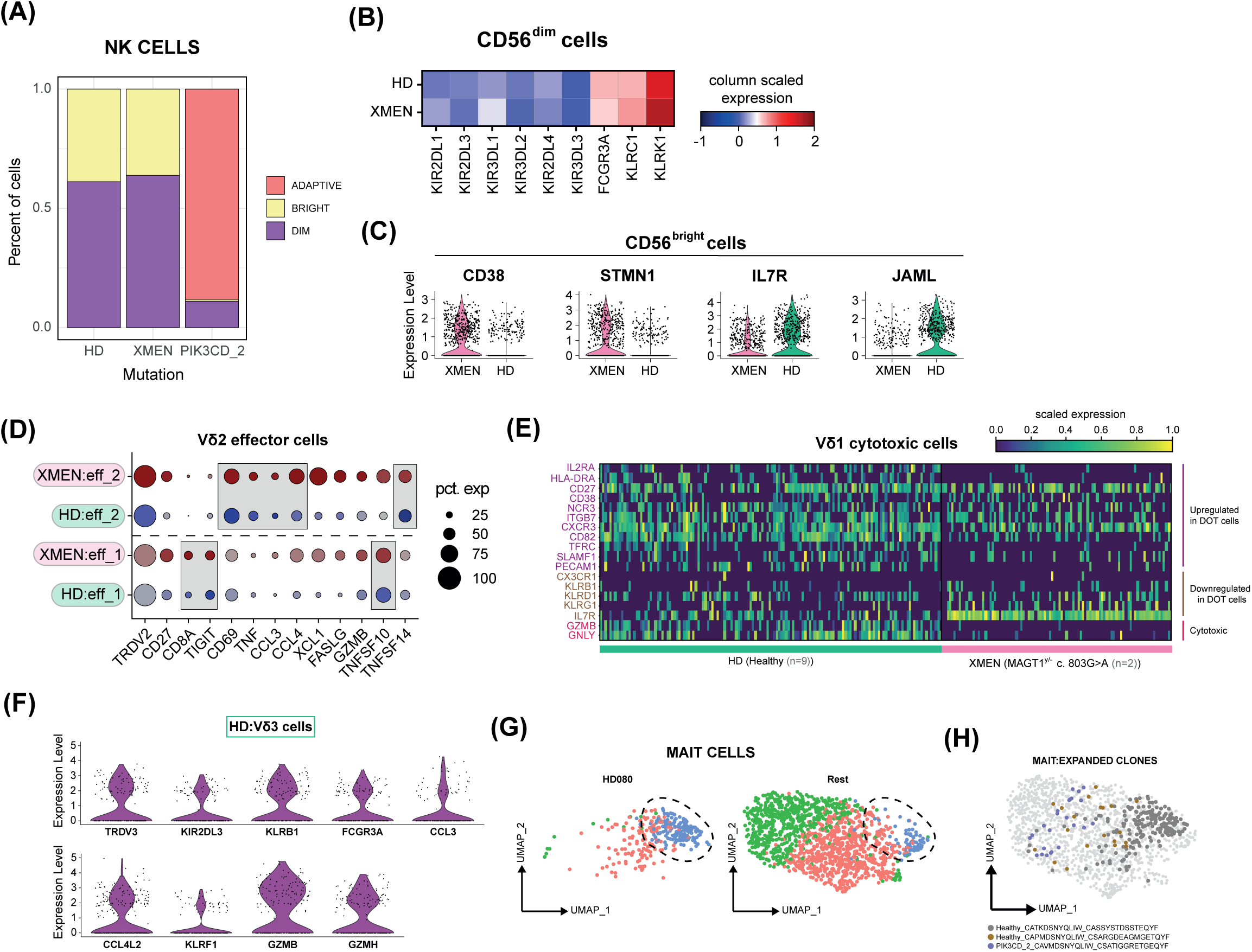
Transcriptomic analysis of innate cells activated against EBV. (A) Stacked bar-plot showing the proportion of dim, bright, and adaptive NK cells per mutation. (B) Matrix plot of *KIR*, *FCGR3A*, *KLRC1*, and *KLRK1* detection per mutation within C56^dim^ cells. (C) Violin plots of some differentially expressed markers between HD and XMEN in CD56^bright^ cells. (D) Split dot plot showing genes upregulated in effector populations of Vδ2 per mutation. Highly coincident events are highlighted in boxes. (E) Scaled heatmap of markers upregulated or downregulated in DOT cells according to Almeida *et al*^26^ within Vδ1 cytotoxic cells per mutation. *GZMB* and *GNLY* expression are also represented. (F) Violin plots of markers expressed in Vδ3 cells. (G) UMAPs showing the distribution of MAIT cells in a healthy donor (HD080) sample and the rest of samples. The active subcluster is highlighted. (K) Representative examples of the most expanded MAIT clones in the UMAP 2D space.

Like CD8^+^ αβ T cells and NK cells, gamma-delta T cells were really only detected in healthy subjects and XMEN patients. Two subpopulations of Vδ2 effector cells that were similar in healthy donors and patients, expressing *CD8A* and *TIGIT*, or classical molecules like *TNF*, *CCL3*, and *CCL4* were found (**Figure 5D, Supplementary Figure 7A**,B). With regard to other gamma-delta T cells, Vδ1 cells expressing homeostatic markers (*CX3CR1*, *KLRG1*, or *GZMK*) were detected in XMEN patients (**Supplementary Figure 7C**-E). Active Vδ1 cells resembling the cytotoxic Delta One T (DOT) cells described by Almeida *et al*^26^ were detected in healthy subjects and XMEN patients, however the Vδ1 cells from XMEN patients completely lacked key cytotoxic molecules like *GNLY* and *GZMB* (**Figure 5E, Supplementary Figure 7F**). Non-conventional Vδ3 effector cells expressing chemokines (*CCL3*, *CCL4L2*), innate molecules (*KLRB1*, *FCGR3A*, *KIR2DL3*), and granzymes were only identified in healthy donors (**Figure 5F**). Finally, although a marked clonal expansion of MAIT cells was detected in one healthy donor, with evidence of some recently described active MAIT identities^27^ (**Figure 5H,I, Supplementary Figure 7G**-K), generally MAIT cells present in peripheral blood do not seem to be essential participants in the cellular immune response to EBV infection.

### 4. Validation of immune subpopulations that respond to EBV: involvement in cell killing

The previous data identified candidate immune cell populations that proliferated or expressed activation markers in response to EBV infection. To validate the transcriptomics data at the protein level in the healthy population, different panels of antibodies were used in flow cytometry. We characterised mature CD4 and CD8 T cells, together with Vδ1 cells, at day 3 and day 6 after EBV stimulation of PBMCs, since these two time points represented an early stage where the frequency of EBV-infected B cells were rapidly decreasing (**Supplementary Figure 8A**), and a late stage with a mix of active and resting effector cells (as depicted in proliferating-based scRNA-seq). Activated, mature CD4 and CD8 T cells were generally found in great proportions in the majority of donors. CD4 T cells were characterised through the expression of CCR2 (**Figure 6A**), and transient expression of GM-CSF (**Supplementary Figure 8B**), whereas responding CD8 T cells were detected using anti-CXCR3 and anti-CCR5 antibodies (**Figure 6B,C**). Further characterisation of T lymphocytes by spectral cytometry revealed that enriched T lymphocytes acquired multiple activation markers (HLA-DR, CD25, CD38) following stimulation with EBV. Additionally, it was shown that active CD8 T lymphocytes expressed KLRG1 and CD27 simultaneously, as well as upregulated GZMB (**Figure 6D,E, Supplementary Figure 8C**). When the functional capacities (IFN-γ and LAMP1 expression) of the T cells that had expanded after incubation with autologous EBV-infected PBMCs were studied, CCR5-expressing adaptive CD4 and CD8 T cells degranulated and upregulated cytokine production after exposure to autologous LCLs (**Figure 6F**, **Supplementary Figure 8D**).

**Figure 6.**
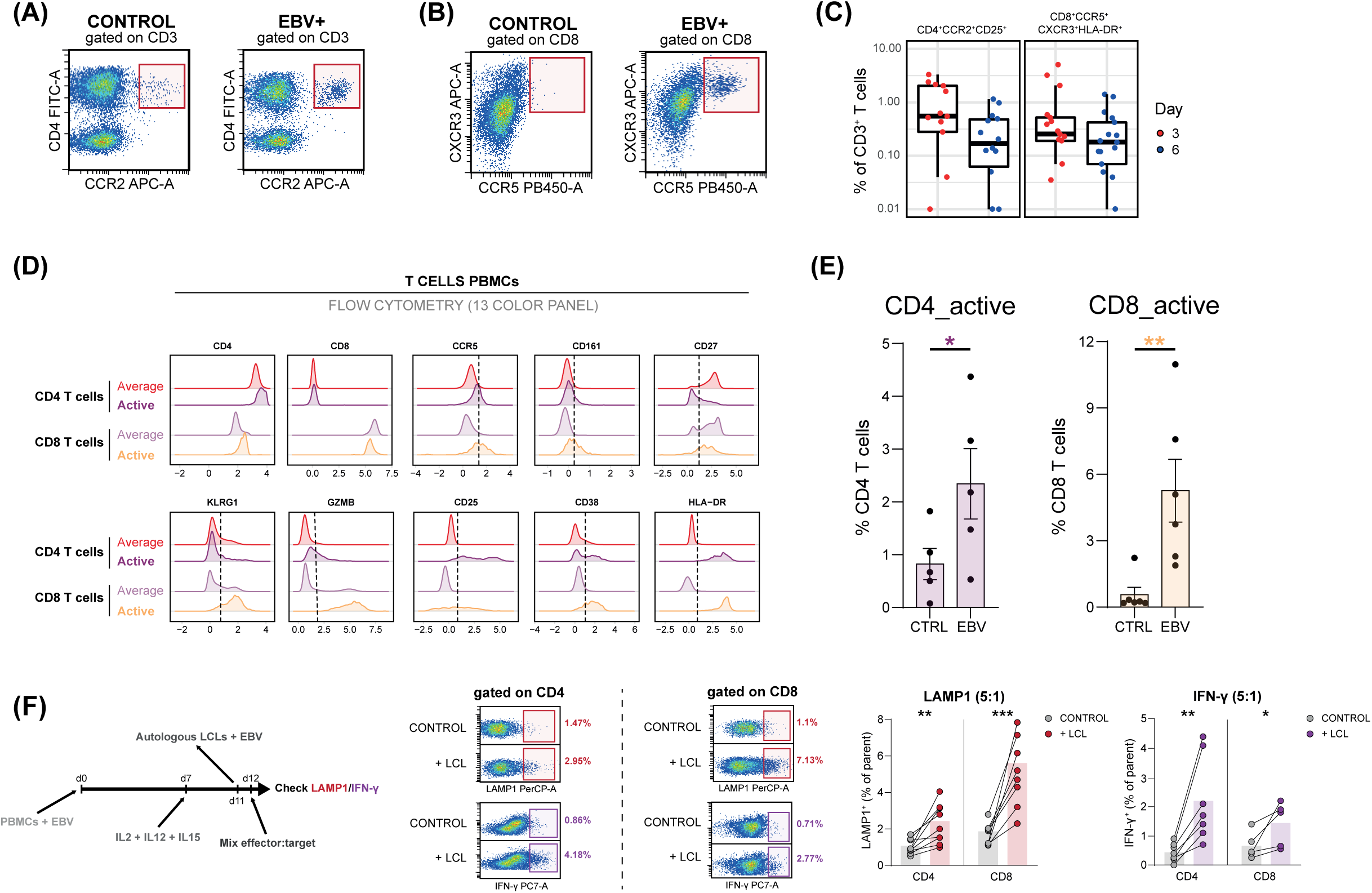
FACS validation of responding cells (I). (A) Representative FACS dot-plots of CCR2-expressing CD4 T cells and (B) CCR5/CXCR3-expressing CD8 T cells in control and EBV-stimulated conditions. (C) Box-plots showing the presence of the selected populations at day 3 and 6 after incubation with EBV. (D) Histograms representing the expression of markers in the average populations and in active clusters detected in the spectral flow cytometry T cell panel. (E) Bar-plots showing the differences in the detection of CD4 or CD8 active clusters between the control and the EBV-stimulated conditions. Paired-sample t-test (*p<0.05, **p<0.01). (F) Expression of LAMP1 and IFN-γ within mature CD4 and CD8 T cells after mixing EBV + cytokine expanded PBMCs with EBV-pulsed LCLs in a 5:1 ratio.

Marked evidence of CD69 upregulation on the innate-like population of Vδ1 cells was found for some donors (**Figure 7A**). The Vδ1 subpopulation was further confirmed to proliferate, although its expression of CD8 was dim and the NKG2C molecule did not seem to be selective for the responding fraction (**Figure 7B**, **Supplementary Figure 8E**,F). We also detected CD8 positive cells expressing CD16, KIR2DL3, and CD161 after EBV incubation, corresponding to non-Vδ2 gamma-delta T cells according to scRNA-seq (**Supplementary Figure 8G**). Similarly, NKG2D-expressing Vδ2 cells upregulating the activation marker CD69 were found at day 3 after EBV stimulation (**Figure 7C**), in great percentage in some donors, and could be further expanded in the presence of IL-2/12/15 (**Supplementary Figure 8H**).

**Figure 7.**
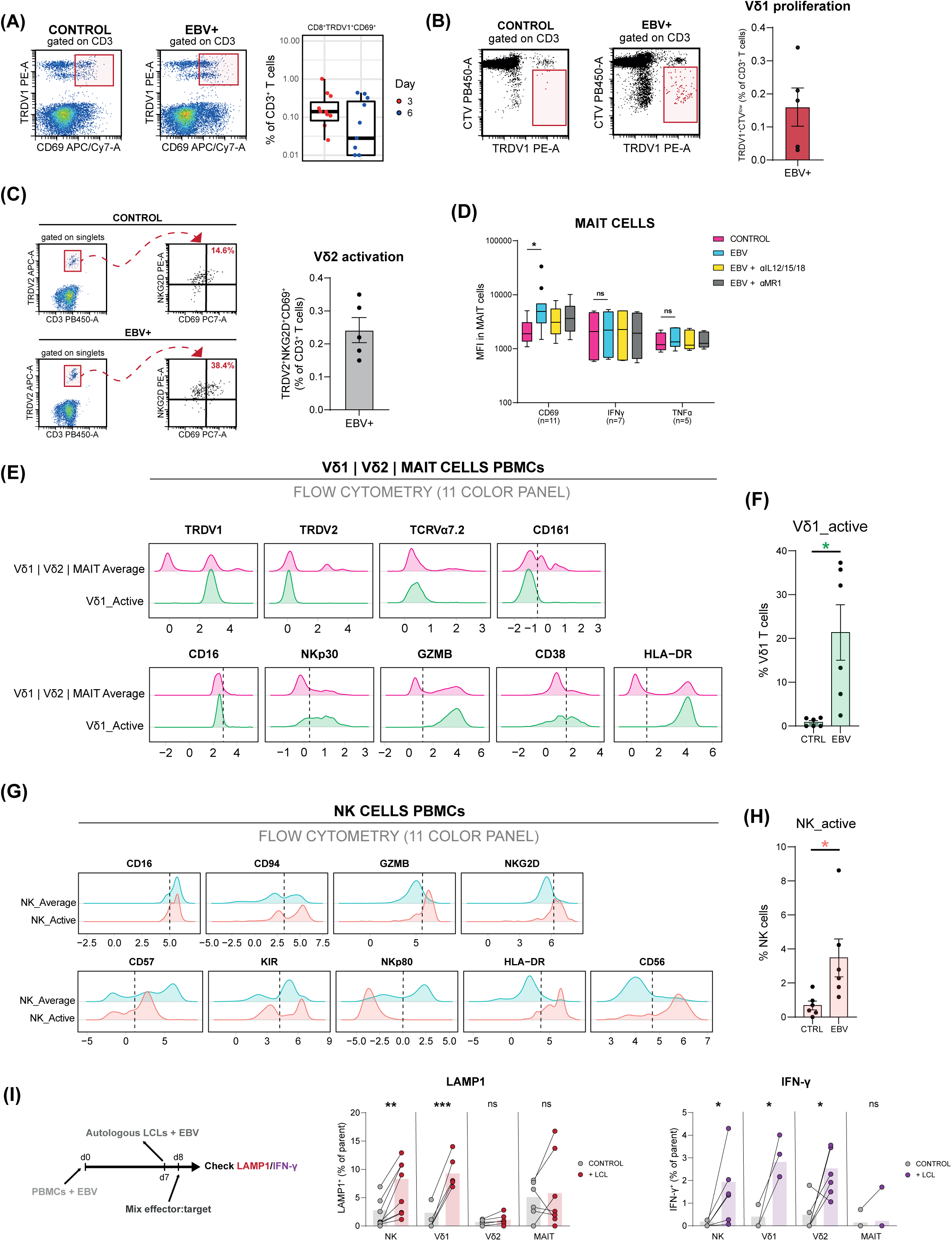
FACS validation of responding cells (II). (A) Representative FACS dot-plot of CD69-expressing Vδ1 cells in control and EBV-stimulated conditions. Box-plot showing the presence of the selected populations at day 3 and 6 after incubation with EBV. (B) CTV dilution within Vδ1 cells after stimulation with EBV at day 6. (C) Upregulation of CD69 in Vδ2 cells after incubation with EBV at day 3. (D) Box-plots indicating the median fluorescence intensity (MFI) of CD69, IFN-γ, and TNF-α within MAIT cells in control and EBV-stimulated conditions, including the blocking with α-IL-12/15/18 (10, 5, and 10 μg/mL) and α-MR1 (2 μg/mL). Outliers are represented as black dots. Paired-sample t-test (*p<0.05). (E) Histograms representing the expression of markers in the average population and in the active Vδ1 cluster detected in the spectral flow cytometry Vδ1 | Vδ2 | MAIT panel. (F) Bar-plot showing the differences in the detection of the active Vδ1 cluster between the control and the EBV-stimulated conditions. Paired-sample t-test (*p<0.05). (G) Histograms representing the expression of markers in the average NK population and in the active NK cluster detected in the spectral flow cytometry NK cell panel. (H) Bar-plot showing the differences in the detection of the active NK cluster between the control and the EBV-stimulated conditions. Paired-sample t-test (*p<0.05). (I) Expression of LAMP1 and IFN-γ within innate populations after mixing EBV-stimulated PBMCs with EBV-pulsed LCLs in a 5:1 ratio.

We further explored the possible activation of MAIT cells after EBV stimulation of PBMCs, but MAIT proliferation at day 6 was generally absent or very low and given its low proportion in blood it is difficult to evaluate the significance of this observation (**Supplementary Figure 8I**). However, we found that in some donors, CD69 upregulation could be significant at day 3, with a weak trend for increased TNF-α production, and this could be partially reversed using blocking antibodies against IL- 12/15/18 or, to a lesser extent, against MR1 (**Figure 7D**).

A combinatorial panel of spectral cytometry allowed for a better understanding of the possible activation of these cell types at day 6, demonstrating, at least with the selection of markers used, that the magnitude of activation of Vδ1 is more substantial than that seen for Vδ2 or MAIT cells, upregulating both activation markers (HLA-DR, CD38) and cytotoxicity molecules (NKp30, GZMB) (**Figure 7E,F, Supplementary Figure 8J**).

Previous work has shown that NK cells are an important lymphocyte population that responds against EBV or LCLs^13^. For this reason, a panel of antibodies was used to visualise changes in NK markers between control and EBV-infected PBMCs at day 6 (**Supplementary Figure 8K**). Interestingly, enriched NK cells upregulated NKG2D or CD94, while the expression of CD57 or KIR was slightly lower (**Figure 7G,H**).

The functionality of these different innate cell populations was directly studied at day 8 after EBV stimulus. In these experiments, incubation with autologous LCLs consistently induced LAMP1 and IFN-γ expression by NK and Vδ1 cells, while Vδ2 only expressed IFN-γ, and MAIT cells did not show any increase of these molecules (**Figure 7I**).

### 5. Focus on Vδ1 cells as a possible adoptive therapy to treat EBV malignancies

Vδ1 cells were identified in both proliferating and activation-based scRNA-seq experiments, expanded considerably in some donors after EBV stimulation (**Supplementary Figure 9A**), and were shown to both degranulate and produce interferon gamma on exposure to EBV-infected B cells. To analyse the features of this cell population (quite infrequent in blood) in more detail, we studied the transcriptome of the EBV-reactive Vδ1 cells detected in healthy individuals. We identified 2 subclusters that formed a gradient of differentiation that culminated in SC2 (**Figure 8A**). Differentiated cells expressed the transcription factors *ZNF683* (Hobit), *IKZF2* (Helios), *ID3*, or *HOPX*, the naive/central memory markers *TCF7* and *LEF1*, the innate markers *XCL1*, *XCL2*, *KLRC2*, *KLRD1*, and *NCR3*, as well as the activation molecules *TNFRSF4* (OX40), *TNFRSF18* (GITR), and *CXCR3* (**Supplementary Figure 9B**). When the transcriptomic differences between our SC1 and SC2 were compared with the protein differences described by Almeida *et al*^26^ between basal Vδ1 cells and their cytotoxic Delta One T (DOT) cells, many coincidences between Vδ1_basal:SC1 and DOT:SC2 were detected (**Figure 8B**). Given these similarities, DOT cells were directly expanded *in vitro* from PBMCs, using the protocol described by Almeida *et al*^26^, obtaining a mean efficiency of 40.53 ± 7.45% (**Supplementary Figure 9C**). In the expansion culture medium, rich in cytokines, and termed “non-resting” condition, a significant degranulation capacity could be noticed after incubation with autologous and heterologous LCLs, super-infected, or not, with EBV (**Figure 8C**). However, given the considerable basal signal for LAMP1 detected in the negative control, subsequent assays were performed after overnight culture of the DOT cells in cytokine-depleted culture medium, termed “resting” condition, in which we were able to clearly distinguish negative and positive values of LAMP1, and in which target cell lysis was evident at effector:target ratios of 5:1 and especially at 10:1 (**Supplementary Figure 9D**,E).

**Figure 8.**
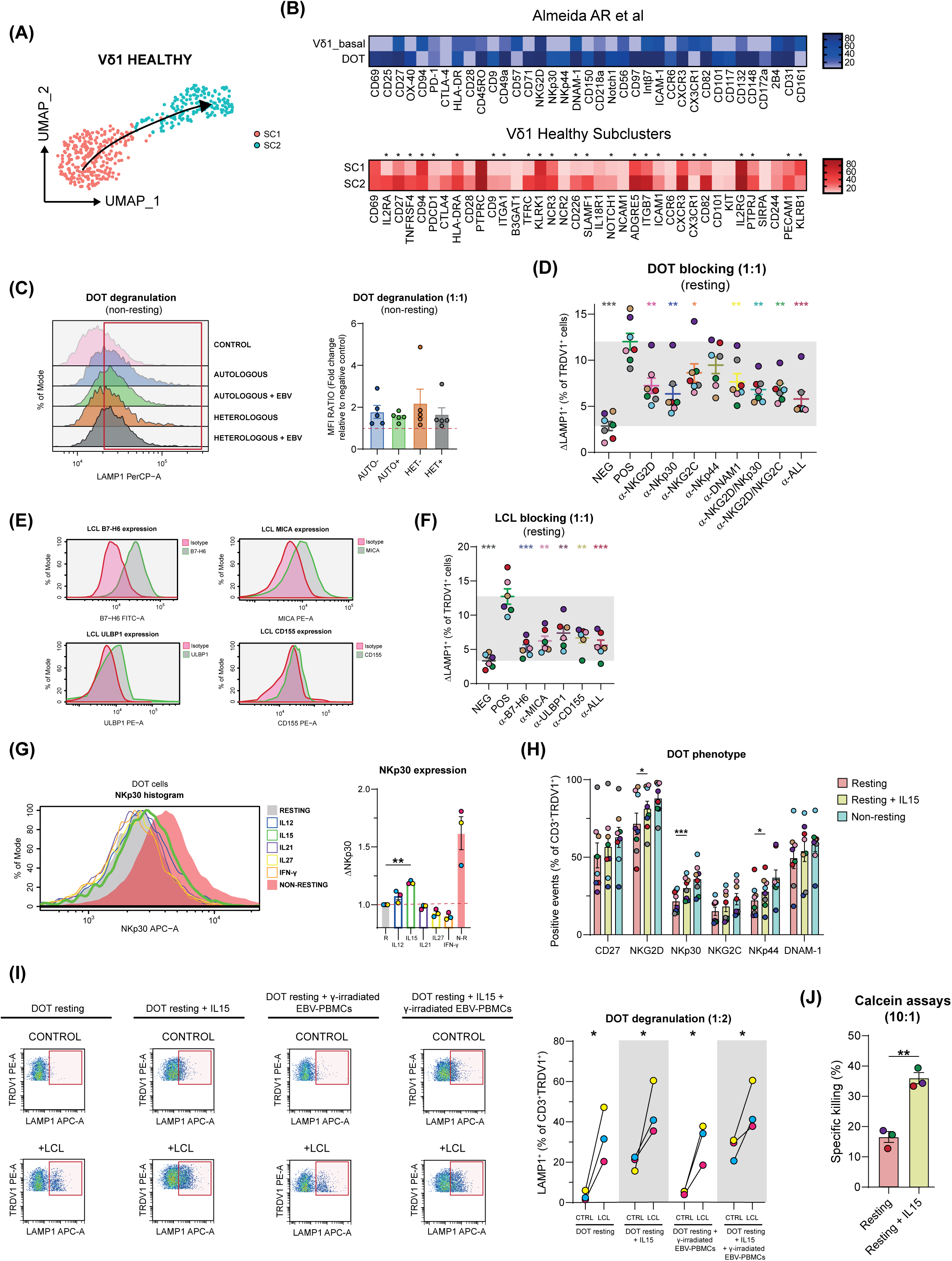
Characterisation of *in vitro* expanded Vδ1 cells. (A) UMAP of healthy Vδ1 cells detected in the activation-based scRNA-seq. Pseudotime trajectory is represented. (B) Heatmap of protein expression of basal Vδ1 and cytotoxic DOT cells described by Almeida *et al*^26^ in blue scale. Heatmap of RNA expression of the genes that encoded the same proteins for SC1 and SC2 in red scale. Coincidences are highlighted with an asterisk. (C) Upregulation of LAMP1 within DOT cells in the non- resting condition when incubating these cells with autologous or heterologous LCLs, pulsed or not pulsed with EBV. (D) Dot-plot of LAMP1 degranulation in the resting condition after using blocking antibodies in DOT cells against NKG2D, NKp30, NKG2C, NKp44, and/or DNAM-1. Paired-sample t-test between positive and the other conditions (*p<0.05, **p<0.01, ***p<0.001). (E) Examples of histograms representing the upregulation of B7-H6, MICA, ULBP1, and CD155 molecules in LCLs. (F) Dot-plot of LAMP1 degranulation in the resting condition after using blocking antibodies in LCLs against B7-H6, MICA, ULBP1, and/or CD155. Paired-sample t-test between positive and the other conditions (**p<0.01, ***p<0.001). (G) Upregulation of NKp30 in DOT cells after incubation with low doses of IL-12, IL-15, IL-21, IL-27, or IFN-γ for 24 hours. Paired-sample t-test (**p<0.01). (H) Bar-plot quantifying expanded DOT cells expressing CD27, NKG2D, NKp30, NKG2C, NKp44, and DNAM-1 in the resting, resting + IL-15, and non-resting conditions. Paired-sample t-test between resting and resting + IL-15 conditions (*p<0.05, **p<0.01, ***p<0.001). (I) Upregulation of LAMP1 within DOT cells in the resting or resting + IL-15 conditions, adding or not gamma- irradiated EBV-stimulated PBMCs, when incubating with LCLs. Paired-sample t-test (*p<0.05). (J) Specific killing of LCLs in resting and resting + IL-15 conditions when using expanded DOT cells at a 10:1 ratio, measured through the release of calcein molecule. Paired-sample t-test (**p<0.01).

We next explored the possible mechanisms by which Vδ1 cells recognised autologous LCLs to trigger LAMP1 upregulation using blocking antibodies specific for NKG2D, NKp30, NKG2C, NKp44, and/or DNAM-1 (**Figure 8D, Supplementary Figure 9F**). In this experiment, the combination of all blocking antibodies diminished the signal of LAMP1 to levels comparable to unstimulated cells, and the NKp30 molecule seemed to be especially significant. Importantly, when we blocked the TRDV1 receptor, we could see that there were no differences in LAMP1 intensity compared to the non-blocked condition, demonstrating that the cytotoxicity of Vδ1 depends on interactions involving multiple innate receptors (**Supplementary Figure 9G**).

Consistent with these data, it was found that LCLs expressed ligands for these innate receptors, especially the NKp30 ligand B7-H6, the NKG2D ligands MICA and ULBP1, and the DNAM-1 ligand CD155 (**Figure 8E, Supplementary Figure 9H**). Regarding this, cell lines showing a latency III phenotype of EBV infection also showed significant expression of B7-H6 and MICA (**Supplementary Figure 9I**). Blocking these molecules on LCLs was sufficient to markedly reduce the degranulation of Vδ1 effectors, although once again, these experiments suggested an important role for NKp30, as the greatest inhibition was achieved when using an anti-B7-H6 monoclonal (**Figure 8F**, **Supplementary Figure 9J**).

Given the apparent importance of NKp30, the upregulation of this molecule in DOT cells was studied after exposure to low concentrations of cytokines (1 ng/mL), and low doses of IL-15 were observed to increase the expression of this receptor, although other receptors like NKG2D or NKp44 also slightly increased (**Figure 8G,H**, **Supplementary Figure 9K**). A degranulation experiment was conducted under conditions with or without IL-15, stimulating or not with gamma-irradiated EBV-stimulated PBMCs. It was observed that when stimulating DOT cells with IL-15, higher degranulation values were obtained against LCLs, but also a higher background signal, while the further addition of EBV-infected PBMCs did not result in any functional change (**Figure 8I**). Nevertheless, it was clear in cytotoxicity assays that IL-15 treatment increased DOT cell killing of autologous LCLs two-fold compared to the condition without IL-15 (**Figure 8J**).

These results indicating that *in vitro* expanded Vδ1 cells can kill EBV-infected cells through innate interactions led us to explore the activity of Vδ1 T cells against LCLs *in vivo*.

### 6. Transferred Vδ1 T cells accumulate in tumour tissue and differentiate towards an effector memory phenotype

For these experiments, nonobese diabetic–scid gamma_c_^−/−^ (NSG) mice were subcutaneously implanted with 1 million LCLs, and 2 days after tumour engraftment, 2 million DOT cells, that had been expanded for 21 days *in vitro*, were intravenously injected into each mouse (**Figure 9A, Supplementary Figure 10A**-C). On sacrifice at day 30 after tumour challenge, low frequencies of human CD45^+^ cells were found in every compartment analysed, but these cells were especially enriched in the tumour tissue, and most of these cells expressed TRDV1 (**Figure 9B,C**). Compared to the Vδ1 cells in blood and spleen, Vδ1 cells isolated from the tumour had markedly upregulated expression of activation markers such as CD39 and CD69, and also tended to polarise towards an effector memory phenotype (CD62L^-^CD45RA^-^) (**Figure 9D, Supplementary Figure 10D**). Moreover, tumour infiltrating Vδ1 cells strongly expressed PD-1 and NKG2A, showed some increase in the expression of CD25, CD56, or NKp30, and contributed overall to a trend of decreasing tumour volume (**Supplementary Figure 10E**).

**Figure 9.**
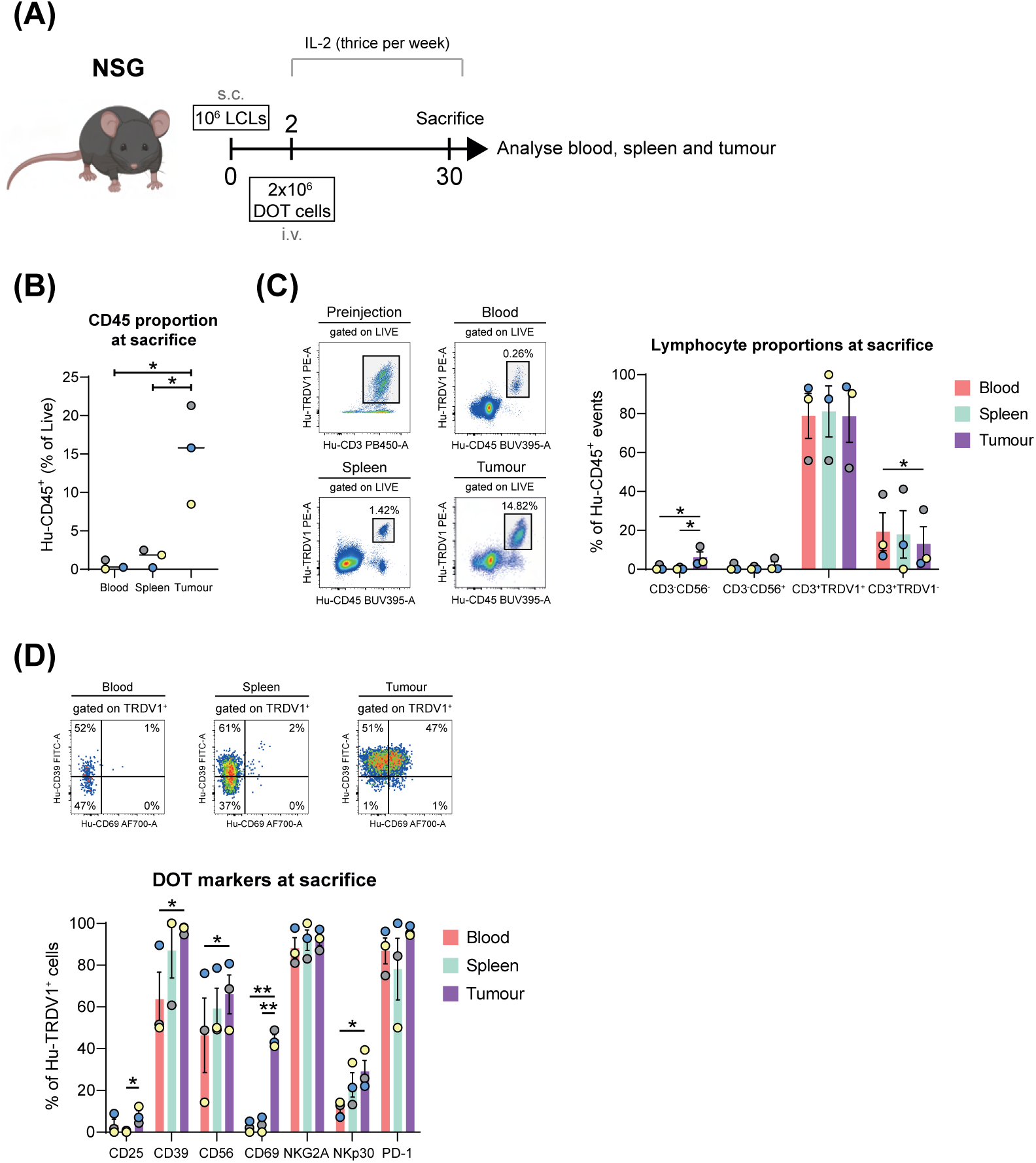
Mice experiment. (A) Schematic of the *in vivo* experiment. (B) Proportion of human CD45^+^ events in blood, spleen, and tumour at sacrifice. Median values are highlighted. Paired sample t-test (*p<0.05). (C) Representative flow cytometry plots and quantification of TRDV1^+^ cells and other lymphocytes in blood, spleen, and tumour at sacrifice. Paired sample t-test (*p<0.05). (D) Representative flow cytometry plots of DOT cells expressing CD39 and/or CD69 in blood, spleen, and tumour at sacrifice and quantification of different molecules included in the flow cytometry panel. Paired sample t-test (*p<0.05, **p<0.01).

## DISCUSSION

In this work, a simple protocol was established which we have used to characterise, for the first time at high resolution, the immune cells present in the peripheral blood of healthy seropositive subjects that comprise the cellular immune response that controls EBV replication *in vivo*. In general, cell-mediated cytotoxicity seems to be critical for effective immune surveillance of EBV and multiple lymphocyte populations contribute to this lytic activity. Secondly, this cytotoxic response is balanced by the activation of CD4 T_reg_ cells. Finally, γδ T cells expressing the Vδ1 T cell receptor use a number of activating NK cell receptors to recognise and kill EBV-infected and transformed lymphoblastoid cells and so are interesting candidates for use in allogeneic cell therapy of EBV-associated lymphoproliferative disorders.

CD4 T cells that proliferated and upregulated activation markers after exposure to EBV were readily detected in healthy donors and in XMEN and XIAP_1 patients, although generally the magnitude of the response was greater in samples from healthy subjects. Cytotoxic CD8 T cell, γδ T cell and NK cell activation could also be detected in samples from healthy subjects and XMEN patients, but not from individuals with mutations affecting XIAP and consistent with this observation, while it was possible to generate CTL lines from healthy donors and XMEN patients, CTLs could not be grown from the XLP-2 patient (XIAP_1). It should be pointed out that although the CTLs from XMEN patients could recognise EBV-infected LCLs and degranulate, the cytotoxic capacity of these cells was much less than for CTLs from healthy donors. Finally, although cellular immune responses to EBV could be detected in both patients and controls, PBMCs from neither XMEN nor XLP-2 patients could restrict the outgrowth of LCLs after EBV infection *in vitro*.

Thus, these data demonstrated that although some lymphocytes from the IEI patients could recognise EBV-infected cells, they were unable to effectively control virus-infected cells. These observations suggested that detailed comparison of EBV-reactive cells from healthy subjects and IEI patients could provide insights into key mechanisms of EBV control. Therefore, we carried out scRNA-seq experiments, comparing cells from healthy donors and IEI patients that either proliferated or expressed activation markers after exposure to autologous EBV-infected B cells. Importantly, for healthy subjects, the immune cell populations that participated in the elimination of EBV-infected cells were consistently detected in both types of experiment of scRNA-seq (proliferation or induced expression of activation markers).

CD8^+^ TCRαβ T cells are generally considered to be key components of the cellular immune response to EBV infection^3,9^ and were prominent among the cells responding to EBV detected in these experiments. In our hands, EBV-reactive cytotoxic T cells expressed *CD27*, important for the expansion and effector function of lytic EBV antigen-specific CD8 T cells^28^, as well as *EOMES*, upregulated in CD8 cells in experiments with EBV tetramers in paired samples from infectious mononucleosis and convalescent patients^29^, and *KLRG1*, a marker that has been found in CD8 T cells undergoing persistent antigen stimulation, as occurs in chronic viral infections (HIV, CMV, or EBV)^30,31^. These cells also upregulated *TIGIT*, a candidate checkpoint receptor that strikingly identified CD8 T cells with increased cytotoxic capacity in HIV studies^32^. In fact, a very high expression of *GZMB* was noted, the classical granzyme that cleaves caspase-3 and caspase-7 in infected or tumour cells^33^. CD8 T cells able to recognise EBV-infected LCLs and degranulate were also detected in XMEN patients, however, as previously reported, the *in vitro*-expanded CTLs from XMEN patients exerted much lower levels of cytotoxicity than CTLs from healthy donors^34^. Some of this is likely due to the much lower levels of NKG2D expression in these patients, but in our experiments we also noted markedly lower levels of expression of genes like *GNLY* and *GZMB*, that are important effectors involved in the cytotoxic function of immune cells. This latter observation differs from previous studies^34^, and this difference may be due to the different stimulation regimes used in the experiments. The CD8^+^ cells here were analysed 6 days after stimulation by exposure to autologous EBV-infected B cells in PBMCs, whereas Chaigne-Delalande *et al* generated CTLs by multiple rounds of stimulation with EBV-LCLs and high dose IL-2. Perforin and granzyme genes are induced during T cell activation and are highest in differentiated memory CD8 T cells^35,36^, moreover IL-2 is known to induce perforin and granzyme expression directly^37^.

The reasons underlying the differences in detection of CD8-expressing CTLs between the scRNA-seq analysis of the XIAP and PIK3CD patients and healthy donors are not clear. Prior work has shown that T-, B- or NK-cell lymphopenia is not frequent in XIAP-deficient patients^38^. Similarly, although PIK3CD patients often present with a progressive CD4 T cell lymphopenia, CD8 T cell counts are generally normal or even high^39,40^. It is possible that the increased propensity of XIAP- and PIK3CD-deficient T cells to enter apoptosis may underlie the lack of CTLs in the cellular immune response to EBV infection^38,40,41^, but whatever the mechanism the association of deficient cytotoxicity in XMEN patients and the absence of CTLs in XIAP and PIK3CD-deficiency strongly support the suggestion that efficient lymphocyte cytotoxicity is very important for control of EBV replication.

CD4 T cells responding to EBV were enriched in a proliferative population expressing *CCR2*, *CXCR3*, *ICOS*, or *GNLY*, showing a Th1-like profile with the highest detected clonal expansion among all CD4 subtypes, and potentially representing a subset of memory effector CD4 T cells with a strong migration and recall capacity^42^. Many *in vitro* isolated EBV-specific CD4 T cell clones have direct cytotoxic capacity against HLA-matched LCL targets and so these data, from short term cultures, suggest that the presence of cytotoxic CD4 T cells in the response to EBV is a genuine feature of this response and not an artifact of the prolonged *in vitro* culture needed to generate human CD4^+^ clones. Analysis of the non-naive cells recovered from the XIAP_2 patient revealed a majority of *IL7R*^+^*KLRB1*^+^*CXCR3*^+^*GZMK*^+^ CD4 T cells. *CXCR3* expression suggested a marked migration capacity and Th1 polarisation, and this population was unique compared to the rest of the detected CD4 T cells as it expressed *GZMA* and *GZMK*, but not *GZMB*, a profile previously identified in cytotoxic CD4 T cells responding to Vaccinia virus vaccination^43,44^. No NK cell response to EBV, and only a very low number of non-naive CD8 T cells was found in XIAP patients, so perhaps these CD4 T cells may represent an attempt to compensate the otherwise poor cytotoxic response to EBV-infected cells.

A significant presence of *FOXP3*-expressing regulatory CD4 T cells was also detected in these experiments, many of them expressing molecules like *CD82*, *HLA-DRB1*, or importantly, *CCR4*, which is known to be upregulated in regulatory T cells with potent suppressive activity^45^. It is interesting to note that lower frequencies of CD25^hi^ regulatory T cells are observed in the peripheral blood of individuals with IM^20^. Perhaps reduced control over the effector response, contributes to the exaggerated CD8 T cell responses that appear to underlie the development of this immunopathological disease. Moreover, since during T cell responses to nonself antigens T_regs_ appear to be important regulators of CD8 T cell priming as well as playing key roles in the induction of strong primary responses and effective memory^46^, it is at least conceivable that the markedly diminished T_reg_ response of the IEI patients may contribute to the impaired cytotoxic responses in these individuals.

Multiple studies, both *in vitro* and *in vivo*, suggest that NK cells play a role in anti-EBV immunity^11,12,13,47^ and in our experiments, significant numbers of NK cells from healthy donors responded to EBV infection. Phenotypically these NK cells were reminiscent of the previously described “early differentiated” NK cells reported to preferentially degranulate and proliferate on exposure to EBV-infected B cells expressing lytic antigens^13^ insofar as these CD56^dim^ cells expressed *KIR* genes at low levels, with moderate expression of *FCGR3A* and *KLRC1*, and high expression of *KLRK1* (mRNA). Intriguingly, XMEN-derived NK cells were somewhat similar to NK cells from healthy donors, although the KLRK1 protein (NKG2D), which is critical for NK cell effector activity against EBV, is not expressed at the cell surface of lymphocytes from these patients. Transcripts for some genes of NK cell function (e.g., *CXCR4*, *IL7R*, *JAML*, *XCL2*), important in the recruitment of other effector cells^48^, were downmodulated in the patient samples. Interestingly, however, unlike CD8 T cells, no marked differences in *GZMB* expression were noted when the transcriptomes of EBV-responding NK cells from healthy donors and XMEN patients were compared. This may simply reflect that essentially all NK cells express *GZMB*, independently of their activation status^35^. Consistent with the paucity of CD8^+^ CTLs in XIAP deficiency, very few NK cells were detected among the EBV-reactive cells. However, different results were obtained in the analysis of two patients with gain of function mutations affecting *PIK3CD*. In PIK3CD_1, around half of the activated cells corresponded to ZNF683-expressing oligoclonal CD8αβ T cells, indicative of tissue resident memory T cells (T_RM_) with long-lived effector capacity^49^. This subpopulation might represent aberrant CD8 T cells, highly cytotoxic but expressing both the *KLRC1* (NKG2A) and *KLRC2* (NKG2C), inhibitory and activating receptors respectively, that interact with HLA-E^50^. In fact, NKG2A-expressing CD8 T cells are thought to appear after repeated antigen stimulation and might represent an exhaustion state^51^. The significance of the oligoclonality of the CD8 CTL expansion is unknown, since analysis of these TCR sequences did not reveal any clues as to the specificity of these receptors (data not shown). On the other hand, the EBV-reactive cells of PIK3CD_2 contained a significant population of adaptive NK cells, that were not detected in samples from other healthy subjects or IEI patients. These NK cells expressed *KLRC2*, *FCGR3A*, *GZMH*, *TIGIT*, and HLA-II molecules, and lacked *FCER1G*, *KLRC1*, *KLRB1*, and *NCR3*, and so clearly represent a phenotype of activated NK cells with adaptive features, typically associated with HCMV infection^52^. Indeed, clinical data confirmed that PIK3CD_1 is seronegative for HCMV, while PIK3CD_2 is seropositive. The significance of these observations is unclear. It is possible that the CD8 T cells expressing innate and lectin-like molecules in PIK3CD_1 might represent some kind of compensatory response, but although EBV infection can alter the repertoire of NK cells in peripheral blood, NKG2C^hi^ adaptive NK cells like those found in PIK3CD_2 are not known to expand in response to EBV infection^53^.

In humans, the innate lymphocyte response to EBV infection has been reported to take two forms. Roughly 50% of donors respond exclusively via NK cells, whereas in the other half of the population both NK cells and Vγ9Vδ2 T cells respond to EBV^14^. In our hands, NK cells responded robustly to autologous LCLs by degranulation and cytokine production. In contrast, LCL-stimulated Vδ2 cells produced IFN-γ, as well as other molecules like *CCL4* or *TNF*, but did not mobilise LAMP1 to the cell surface, suggesting a more “helper-like” role for these gamma-delta Vδ2 T cells, although cell killing might be modulated through other pathways not involving LAMP1, e.g., *FASLG* and several members of the TNF and TNFR superfamilies (data not shown).

Strikingly, however, in the majority of donors tested, Vδ1 cells were also found to upregulate CD69 and, importantly, since these cells are normally thought of as tissue-resident and CD69 expression could be just intrinsic^54^, to proliferate. These cells could greatly upregulate NKp30 and GZMB, molecules with an evident cytotoxic function. To a lesser extent, T cells expressing CD16, KIR2DL3, and CD161, that might represent non-Vδ2 gamma-delta cells, especially a fraction enriched in TRDV3^+^ cells according to transcriptomic analysis, also upregulated the expression of activation markers.

The detection of Vδ1/Vδ3 cells was intriguing because these cells are primarily found in the intestinal mucosa, but consistent expression of previously described activation markers was noticed, including *NCR3*, a molecule that appears to be key for the cytotoxic function of this cellular subtype^55^. Further, Vδ1 T cells that expanded in our *in vitro* system responded robustly to autologous EBV LCLs by degranulation and IFN-γ production. Unlike the better characterised Vγ9Vδ2 gamma-delta cells, our data are consistent with prior clinical observations showing that Vδ1 T cells can expand *in vivo* during EBV-induced infectious mononucleosis^56^ and upon EBV infection after cord blood transplantation^57^. Vδ1 T cells also expand *in vitro* in response to ligands expressed by EBV-infected Burkitt lymphoma cells and transformed B lymphocytes, and can mediate cytotoxicity against autologous EBV–LCLs^58,59^. Using blocking experiments with monoclonal antibodies, we showed that Vδ1 T cell recognition of the EBV-LCLs was mediated by multiple activating NK receptors expressed by these γδ T cells and we could also confirm the expression of the ligands for these receptors by the latently infected B cells. Strikingly, TCR blockade had no effect on recognition of the target cell. As mentioned previously, analysis of our scRNA-seq data revealed substantial overlap between the EBV-reactive TCR Vδ1 γδ T cells identified in our experiments and the “Delta One” T (DOT) cells described by Almeida *et al*^26^. Importantly, comparison of the Vδ1 expansions in IEI patients with those of healthy donors revealed that the patient Vδ1 T cells lacked expression of key cytotoxic effector molecules like GZMB and granulysin, further reinforcing the idea that cytotoxicity, mediated by multiple effector lymphocyte populations, is critical for immune control of EBV.

DOT cells have been shown to be highly effective in eliminating cancer cells in patients with defects in HLA class I expression^60^ and their use as a cell therapy for a range of cancers is being actively explored^61^. Given the marked similarities between DOT cells and the EBV-reactive TCR Vδ1 γδ T cells identified in our experiments we hypothesised that Vδ1 T cells might be used as a therapy for EBV-associated cancers, particularly in patients with some degree of immunocompromise.

Individuals with weakened immune systems, due either to inborn errors of immunity or immunosuppression, e.g., after hematopoietic stem cell or solid organ transplant, often suffer severe EBV-related complications including infections and lymphoproliferative diseases. EBV infection is also a frequent cause of hemophagocytic lymphohistiocytosis when cell-mediated cytotoxicity is impaired. Apart from Rituximab, a monoclonal antibody against CD20, that targets both EBV-infected and healthy B cells^62^, further weakening immunity, antiviral pharmacotherapy is not available for EBV, and so strategies based on adoptive transfer of EBV-specific T cells have been used as a therapeutic for immune-compromised patients with EBV-associated diseases^63,64,65^. These approximations can be very effective, but the high cost and complexity of manufacturing individual T-cell products limit accessibility, and patient-derived products can show important variations in effectiveness. To avoid these limitations, the use of allogeneic donor-derived banks of characterised HLA-typed EBV-specific T cell lines from normal donors has also been explored^66^. However, here the degree of HLA matching is an important limiting factor, particularly in the setting of rare HLA alleles that are not known to mediate recognition of immunodominant EBV epitopes. Finally, resistance to EBV-specific cellular therapy due to viral escape mutations in peptide epitopes has also been described^67^.

In our experiments, expanded Vδ1 cells recognise EBV-infected B cells via multiple activating NK receptor/ligand interactions to mediate cytotoxicity and cytokine production in a non-MHC/non-TCR dependent fashion. These observations suggest that these lymphocytes could be candidates for an allogeneic, “off-the shelf” cell therapy of EBV-infection that would require the tumour to undergo multiple mutations to become resistant and so we explored the ability of these cells to act against latently-infected EBV LCLs in a xenograft lymphoma model. In this initial study, the transferred Vδ1 T cells persisted in mice for at least 30 days after transfer and specifically accumulated in the tumours. The tumour-infiltrating Vδ1 T cells upregulated CD69 and CD39, among other markers, demonstrating activation/homing against LCLs even at day 30. On the other hand, the transferred Vδ1 T cells that were found in the spleen were of a central memory phenotype, as expected for secondary lymphoid organs. Although there was a trend towards reduced tumour growth in mice that received Vδ1 T cells, this was not statistically significant. This may reflect the low numbers of mice used and perhaps also the expression of the inhibitory receptors PD-1 and NKG2A by Vδ1 T cells *in vivo*. These data suggest that combination with monoclonal antibodies targeting these checkpoint inhibitors might enhance the immune control of tumour growth by the transferred Vδ1 cells. Another limitation of these experiments is that while the effector ability of the expanded Vδ1 cells depended on various activating NK receptors, NKp30 (*NCR3*, CD337) seemed to be particularly important for recognition, and in fact LCLs were enriched in ligands for the NKp30 molecule. The transcriptomic analyses suggested that the more active cells were those with greater upregulation of this receptor. However, NKp30 was expressed by only a small proportion of the Vδ1 cells expanded *in vitro* by the current protocol^26^. Further investigation is required to optimise the culture conditions to maximally upregulate the expression of this marker in the *in vitro* expanded type 1 gamma-delta T cells and, hopefully, augment the potency of the cytotoxic response against EBV-associated tumours.

In conclusion, the use of single-cell phenotypic characterisation technologies in healthy carriers and IEI patients particularly susceptible to EBV disease has allowed, for the first time, a global characterisation of the cell types and subtypes that contribute to efficient immune control of this virus. The sequencing in parallel of healthy seropositive donors and a range of IEI patients has confirmed the role of classical CD8^+^ cytotoxic T cells positive for *CD27* and *EOMES* in the proper elimination of infected cells. However, other effector lymphocytes like NK cells and TCR Vδ1 cells expressing *CXCR3* and *NCR3* also appear to contribute to this cytotoxicity, and may represent a novel approach for therapy of EBV-associated diseases.

## LIMITATIONS

This study has focussed on the characteristics of the immune response to EBV in the peripheral blood of the analysed individuals, and it has not been possible to consider differences that may arise in other physiologically relevant contexts of infection by this virus, such as secondary lymphoid tissues like tonsils. Moreover, although the transcriptome and TCR of single cells from all available IEI subjects have been sequenced, sample-volume limitations have prevented, in many cases, additional experiments, particularly for XIAP_2, PIK3CD_1, and PIK3CD_2. Finally, although the Vδ1 cell-injection experiment in mice has yielded promising results, it is evidently limited by the number of mice used and by the use of a single LCL batch.

## METHODS

### 1. Samples

#### 1.1. Healthy donor samples

Buffy coats of healthy donors were obtained from the Regional Transfusion Centre (Madrid), whereas the samples of IEI patients came from various hospitals. All participants gave their informed consent and the ethical permission and experimental procedures were approved by the bioethics committee of CSIC (078/2019).

#### 1.2. Immunodeficient samples

##### 1.2.1. MAGT1^y/-^

Mutations in *MAGT1* gene comprised X-linked immunodeficiency with magnesium defect, Epstein-Barr virus infection, and neoplasia (XMEN). In this work, we used 3 samples from affected siblings carrying the same mutation in the *MAGT1* gene (c. 803G>A, p. Trp268Ter) (**Supplementay Figure 11A**). These patients shared the following features in peripheral blood: decrease in the expresion of NKG2D in CD8 and NK cells (**Supplementary Figure 11B**), predominancy of B cells with a naive phenotype and correct distribution of intracytoplasmic kappa and lambda light chains, inverted CD4/CD8 ratio (**Supplementary Figure 11C**), and diminished proliferation in response to mitogenic stimulation with anti-CD3. Moreover, each of them had the following clinical particularities:

- XMEN_1: male, 28 years. Non-Hodgkin lymphoma diagnosed at 24 years. Immune thrombocytopenic purpura (ITP). Untreated hypogammaglobulinemia.

- XMEN_2: male, 25 years. Hodgkin lymphoma diagnosed at 18 years. Common variable immunodeficiency treated with immunoglobulins.

- XMEN_3: male, 22 years. Hodgkin lymphoma diagnosed at 5 years. Immune thrombocytopenic purpura (ITP). Intracranial bleeding. Splenectomy.

##### 1.2.2. XIAP^y/-^

Two unrelated XIAP^y/-^ patients were analysed:

- XIAP_1: c. 658C>T, p. His220Tyr; diagnosed with XLP-2. First episode of HLH at 3 years, treated with corticosteroids, cyclosporine, and etoposide. The patient presented pancytopenia, elevated liver enzymes, splenomegaly, and titres of CMV IgM^+^ IgG^+^, VCA IgM^+^, EBNA IgG^-^. In the bone marrow, a significant increase in mature macrophages and monocytes, erythroid hyperplasia, hemophagocytosis, and cannibalism were observed. Lymphocytes appeared atypical, and no intracellular parasites were detected.

- XIAP_2: c. 1452C>G, p. Cys234Trp; diagnosed with XLP-2. Clinical onset at the age of 4 years, following several episodes of HLH treated with corticosteroids and cyclosporine. Hypogammaglobulinemia and slightly decreased B lymphocyte population in blood.

##### 1.2.3. PIK3CD^GOF^

Two unrelated patients with GOF mutations in the *PIK3CD* gene were analysed:

- PIK3CD_1: c. 1570T>G, p. Tyr524Asp; diagnosed with APDS. Clinical onset at the age of 19 years with systemic lymphoproliferation (atypical lymphoproliferation CD3^+^ and CD20^+^, EBV-), EBV serostatus positive, no circulating EBV DNA. CMV serostatus negative. In blood, T cells skewed towards activated/effector phenotype, and expansion of CD19^hi^CD21^low^.

- PIK3CD_2: c. 3061G>A, p. Glu1021Lys; diagnosed with APDS. Clinical onset with recurrent bacterial infections, on IgRT from the age of 16. From the age of 27 years, diffuse lymphoproliferation (atypical lymphoproliferative episode CD20^+^, EBV-). recurrent EBV infection with circulating EBV DNA without complications. CMV serostatus positive. In blood, T cells skewed towards activated/effector phenotype, circulating B cells with the exclusive transitional phenotype, expansion of CD19^hi^CD21^low^. Failure to respond to CpG stimulation.

### 2. *In vitro* assays

#### 2.1. Isolation of PBMCs

Peripheral blood mononuclear cells (PBMCs) were isolated from buffy coats by centrifugation on Ficoll-HyPaque. PBMCs were either used fresh in assays, or cryopreserved.

#### 2.2. Isolation of B95-8 EBV

Stocks of EBV were prepared from the marmoset B-lymphoblastoid cell line (B95-8, ATCC CRL 1612). Cells were harvested in complete RPMI 1640 + 10% FBS, and then were centrifuged and plated in a 75 cm^2^ flask in complete RPMI 1640 + 5% FBS at 10^6^ cells/mL for 96 hours. The cells were then centrifuged at 2500 rpm for 10 minutes, and the EBV-containing supernatant was collected through a 0.45 μm sterile filter.

#### 2.3. Stimulation of PBMCs

PBMCs were generally stimulated using an optimised protocol in which 10^6^ cells were incubated with 1 mL of EBV aliquot (MOI ∼ 5) for 2 hours in complete RPMI 1640 + 10% FBS medium. This time was enough to permit infection of B cells while the rest of cell types maintained their ability to respond. After that, cells were washed 3 times in RPMI and were cultured at 2x10^5^ cells/well in 96-well U bottom plates for either 3 or 6 days. The optimal set of conditions was tested according to the proliferation of CD4, CD8, or NK cell effector lymphocytes after trying different variants of the protocol (**Supplementary Figure 11D**).

For peptide assays, the same protocol was followed, except that lymphocytes were treated with IL-2 (Peprotech, 5 U/mL) at day 7, and then restimulated with a mix of commercial peptides against EBV or HCMV (Miltenyi Biotech) at day 12 during 6 hours in the presence of Brefeldin A (BioLegend, 5 μg/mL).

#### 2.4. Generation of LCLs

Donor-specific lymphoblastoid cell lines (LCLs) were generated from PBMCs in 2 ways:

- PBMCs were exposed to EBV aliquots for 2 hours and cultured in 96-well U bottom plates at very low confluence (10^4^ cells/well).
- PBMCs were pre-incubated with PHA (Thermo Fisher, 5 μg/mL) for 30 minutes, then stimulated with EBV aliquots for 2 hours and cultured together with FK-506 (Merck, 10 μg/mL) in 96-well U bottom plates at high confluence (2x10^5^ cells/well).

The RPMI medium was replaced weekly, and growing LCLs were expanded into 24 and 6 well plates, and finally to 25 cm^2^ flasks, in a process taking about 6 weeks.

#### 2.5. CTL assays

Donor-specific cytotoxic T lymphocytes (CTLs) were generated from PBMCs by multiple rounds of stimulation with γ-irradiated autologous LCLs (40 Gy, ratio 1:40 LCLs/PBMCs) in the presence of IL-2 (Peprotech, 50 U/mL). Specific killing assays were conducted incubating CTLs with autologous or heterologous LCLs and evaluating target cell viability using Annexin V or the NK cell degranulation marker LAMP1 after a period of 4 hours.

#### 2.6. Regression assays

Regression assays were performed to determine the minimum confluence of PBMCs needed to avoid the formation of LCLs per donor and condition when cells are infected with EBV. To this end, monocyte-depleted PBMCs were infected for two hours and plated in serial dilutions from 2x10^5^ to 6x10^3^ cells/well in 96-well U bottom plates during 4 weeks. The appearance of cell clumps was identified as growing LCLs, and strength of the regression was quantitatively expressed as the minimal number of cells required for 50% growth inhibition.

#### 2.7. *In vitro* expansion of DOT cells

Delta One T (DOT) cells were grown directly from PBMCs using a protocol similar to the one described by Almeida *et al*^26^. Specifically, γδ T cells were MACS-isolated using the TCRγ/δ+ T Cell Isolation Kit (Miltenyi Biotec), and cells were cultured in 96-well U bottom plates in serum-free culture medium (OpTmizer-CTS) supplemented with 5% autologous plasma and 2 mmol/L l-glutamine (Thermo Fisher Scientific) during 3 weeks using different combinations of cytokines: 1 week with IL-4 (Immunotools, 100 ng/mL), IFN-γ (Immunotools, 70 ng/mL), anti-CD3 (OKT-3, BioLegend, 70 ng/mL), IL-21 (Peprotech, 7ng/mL), and IL-1β (Peprotech, 15 ng/mL); 1 week with IL-15 (70 ng/mL), IFN-γ (30 ng/mL), and anti-CD3 (1 μg/mL); 1 week with IL- 15 (100 ng/mL), IFN-γ (200 ng/mL), and anti-CD3 (70 ng/mL). These cells were used for evaluating the presence of the degranulation marker LAMP1 after 4 hours of incubation with autologous LCLs, in the presence or not of blocking antibodies against NKG2D (Merck, A10), NKp30 (BioLegend, P30-15), NKG2C (R&D Systems, #134522), NKp44 (BioLegend, P44-8), DNAM-1 (R&D Systems, #102511), B7-H6 (BioLegend, W20205G), MICA (R&D Systems, #159207), ULBP1 (R&D Systems, #170818), and/or CD155 (BioLegend, A18179E), 10 μg/mL each one. Specific killing experiments were assayed through the detection of calcein fluorescence in a Victor Nivo Microplate reader after incubating labelled LCLs with unlabelled DOT cells during 3 hours.

### 3. *In vivo* experiments

#### 3.1. Mice injections

These experiments were carried out using a previously described protocol^68^ with the following modifications: LCL tumours were injected subcutaneously into the left flank of the mice under isoflurane narcosis. 1x10^6^ tumour cells were washed and resuspended in PBS and immediately before injection were mixed at a 1:1 V/V ratio with Corning® Matrigel® Growth Factor Reduced (GFR) Basement Membrane Matrix (Corning).

Two days after tumour implantation, 2x10^6^ adjusted DOT cells were adoptively transferred by tail vein injection. To favour the persistence of transferred cells, mice were injected intraperitoneally (i.p.) with 20 × 10^4^ IU IL-2/mouse every second day. Tumour size was monitored by calipering (3x/week or daily). Tumour volume was calculated using the formula: Length x Width^2^/0.52.

General health was monitored by weighing and health parameter scoring (3x/week or daily, according to animal license). Persistence of adoptively transferred cells was monitored by tail vein bleeding. Blood was treated with ammonium chloride potassium (ACK) lysis bufer (Gibco), washed with PBS, stained, and analysed by flow cytometry. All animal experiments were performed according to an approved license by the veterinary office of the canton of Zurich, Switzerland (ZH067/23).

### 3.2. Isolation of single cell suspensions at sacrifice

At sacrifice, blood was collected by heart puncture and treated with ACK lysis buffer to lyse erythrocytes and isolate PBMCs. To isolate splenocytes, spleens were harvested, meshed through a 70 µm cell strainer and lymphocytes isolated by Ficoll-Paque (GE Healthcare) density gradient centrifugation. To isolate tumour infltrating lymphocytes (TILs), the skin covering the tumour was removed and tumours were cut into small fragments with scissors, digested in DMEM (Life technologies) supplemented with 2% heat-inactivated FBS (Biochrome), 1 mg/mL collagenase IV (Roche), 40 µg/mL DNase I (Roche) and 1.2 mM CaCl_2_ for 45 min at 37°C and 200 rpm. After digestion, tumours were extensively washed in supplemented RPMI to stop the digestion, meshed through a 70 μm cell strainer and washed twice with PBS to remove excess of debris.

All single cell suspensions were then stained for 20 min at 4°C and analysed by flow cytometry.

### 4. Flow cytometry

#### 4.1. Surface and intracellular staining

The complete list of antibodies used in this work is given in **Supplementary Table 3**. For surface staining, cells were washed in PBS containing 1% FBS, 0.5% BSA, and 0.05% sodium azide (PBA), and incubated with a selected panel of antibodies for 30 minutes at 4°C. For intracellular staining, surface-stained cells were washed, fixed in 4% paraformaldehyde, and permeabilised in 0.5% saponin prior to incubation with selected antibodies for 40 minutes at room temperature. Samples were acquired on a CytoFLEX (Beckman Coulter) or an Aurora 5L (Cytek), and analysed using CytExpert v2.5, SpectroFlo v2.2.0.3, or R software. For multiparametric flow cytometry analysis, FCS files exported from manual gating were processed using the R CATALYST package (v1.28.0), applying cofactors ranging from 150 to 3000.

#### 4.2. Cell-sorting

Fluorescence-activated cell sorting (FACS) was used to isolate live PBMCs diluting CTV or expressing CD69 and/or CD25 after EBV stimulation. For this, cells were surface stained using the protocol described in the previous subsection and resuspended in EDTA-containing PBS. Cells were separated by fluorescence using a MoFLO XDP Cell Sorter (Beckman Coulter) or a CytoFLEX SRT Benchtop Cell Sorter (Beckman Coulter). Sorted cells were collected directly in PBS with 0.04% BSA and washed immediately prior to single-cell sequencing.

### 5. scRNA-seq

#### 5.1. Sequencing

PBMCs from healthy or immunodeficient samples were stimulated *in vitro* with EBV for 6 days and isolated using FACS based on proliferation (CTV dilution) or activation (CD69 and/or CD25 expression) of cells within the live gate. Samples from different donors were multiplexed using antibodies directed against beta2- microglobulin. Gel beads-in-emulsion (GEMs) were generated in a Chromium controller (10x Genomics, San Francisco), and Gene Expression (GEX) and hashtag-oligo (HTO) scRNA-seq libraries were constructed using a Chromium Next GEM Single Cell 5’ Kit v2. Reads were sequenced in a paired-end strategy (26bp Read1, 10bp Index1, 10 bp Index 2 and 90bp Read2) on a P3 flow cell (100 cycles) of the NextSeq 2000 (Illumina) at the Genomics Unit of the National Centre for Cardiovascular Research (CNIC, Madrid, Spain).

#### 5.2. Analysis

Cell Ranger (v6.0.2) count was used to align and quantify raw gene expression data coming from GEX and HTO FASTQ files. Samples were aligned against the human reference genome (GRCh38-2020-A), and against unannotated and partially annotated EBV genomes. Gene expression matrices were analysed in R software (v4.0.1).

##### 5.2.1. Data cleaning

High-quality singlets were retained filtering out cells with <200 and >7,500 features, and with a mitochondrial content >25%. Oligo-tagged antibodies were additionally used to discriminate doublets, since hashtag counts were normalised using Centered Log-Ratio tranformation and demultiplexed through HTODemux pipeline.

##### 5.2.2. Clustering

Seurat (v4.0.2) was the main R package used for data manipulation and visualisation. Individual Seurat objects (containing preprocessed data from one 10x run) were merged or integrated and GEX counts were normalised and log-transformed. CellCycleScoring() function was used to avoid strong tendencies of clustering based on proliferation genes. Data was scaled performing regression by cell cycle, mitochondrial gene content and/or sample in order to correct batch effects. Principal component analysis (PCA) was performed on variable features, and the first N principal components, estimated through the ElbowPlot() method, were used as input for Shared Nearest Neighbor (SNN) clustering and UMAP visualisation.

Cell clustering was performed at a resolution of 0.5 or 1.2, and FindMarkers() function was used to explore cluster-specific features and perform a general annotation, that was contrasted mapping the merged Seurat object to the Azimuth human PBMC reference dataset (https://azimuth.hubmapconsortium.org/). To better explore differences between clusters, DotPlot(), VlnPlot(), FeaturePlot(), and DoHeatmap() functions from Seurat R package were used, as well as dittoHeatmap() from dittoSeq (v1.6) R package, and stacked_violin(), dotplot(), matrixplot(), tracksplot(), and correlation_matrix() from Scanpy (v1.9.1) Python toolkit.

##### 5.2.3. Pseudobulk

Pseudobulk differentially expressed genes (DEGs) between conditions were calculated with the Seurat AverageExpression() function, and were represented with pheatmap (v1.0.12) R package, prefiltering out features with <0.1 scaled expression.

Pseudobulk principal component analysis (PCA) was conducted through DESeq2 (v1.36.0) R package, running a differential expression analysis within the whole dataset (except B/plasma cells), and regressing out by condition (Healthy/Immunodeficient). Differences between samples were also explored within the DESeq2 object computing pairwise correlation and representing the results with pheatmap (v.1.0.12).

##### 5.2.4. Cell-cell interaction

CellPhoneDB (v2.1.7) Python package was used to predict ligand-receptor (LR) interactions between clusters with the statistical_analysis() method. Cell clusters with >20 cells per condition were included and the log-scaled counts were used as input with default settings. Significant LR interactions (p <0.05) were included to create representations with CrossTalkeR (v1.3.0) R package. Specifically, the plot_cci() function was employed to exhibit the ligand-receptor networks for each condition, where the edges are weighted by the number of interactions and the sum of weights of the interaction-pairs obtained with CellPhoneDB.

##### 5.2.5. Pseudotime

For pseudotime analysis, Vδ1 cells were subsetted and the Monocle2 (v2.18.0) method was used to build single cell trajectories based on Discriminative Dimensionality Reduction with Trees (DDRTree) algorithm, as described^69^.

### 6. scTCR-seq

#### 6.1. Sequencing

TCR transcripts from proliferation or activation-based sorted PBMCs were obtained using a 10x Genomics human T cell Chromium Single Cell V(D)J Enrichment Kit. Reads were sequenced in a paired-end strategy (26bp Read1, 10bp Index1, 10 bp Index 2 and 90bp Read2) at the Genomics Unit of the National Centre for Cardiovascular Research (CNIC, Madrid, Spain).

#### 6.2. Analysis

Cell Ranger (v6.0.2) pipeline was followed to filter high-confidence alpha and beta TCR chains. Filtered contigs were loaded in R software (v4.0.1) or Python (v3.8.5) and were mainly analysed with the scRepertoire (v1.7.0) and immunarch (v.0.9.0) R package or the Scirpy (v0.10.1) Python toolkit, respectively.

##### 6.2.1. Clonality and chain usage

Clonal expansion was explored after creating a h5ad file from the R Seurat object through the sceasy (v0.0.6) R package. This file was loaded in Python (v.3.8.5) and analysed with Scanpy (v1.8.1) and Scirpy (v0.10.1).

Chain usage was represented in bar graphs using the vizGenes() function of scRepertoire R package (v.1.7.0) or the vdj_usage() function of the Scirpy (v.10.1) Python toolkit. For further analysis, the geneUsageAnalysis() function of the immunarch R package (v.0.9.0) was used to cluster samples (through k-means in a PCA space) according to the differential usage of TRBV chains.

##### 6.2.2. Epitope annotation

The VDJdb database was used to annotate those cells with detection of a TCR that was already annotated to be specific to EBV. For that purpose, the query_annotate() function of the Scirpy (v.10.1) Python toolkit was implemented to filter only EBV matches with no redundancy.

##### 6.2.3. GLIPH

The “Grouping of Lymphocyte Interactions by Paratope Hotspots” (GLIPH) algorithm was used to group similar CDR3βs within same convergence groups. Convergence groups are, per definition, a set of multiple TCRs from one or more individuals that bind the same antigen in a similar manner through similar TCR contacts. The gliph-group-discovery.pl command was executed in bash (v5.0.17) under the default parameters, and outputs were further analysed in R software.

## DATA AVAILABILITY

scRNA-seq data of proliferation and activation experiments are available at Gene Expression Omnibus (GEO) under the accession code GSE277638.

## CODE AVAILABILITY

R code related to the main scRNA-seq analyses and figures can be found at https://github.com/algarji/PBMCs_EBV_scRNA-seq.

## COMPETING INTERESTS

The authors declare no competing interests.

## FUNDING

This work was supported by grants PID2020-115506RB-I00 and PID2023-149574NB-I00 to HTR (Spanish Science agency/European Regional Development Fund, European Union, AEI/FEDER, EU). LIGG is supported by the Instituto de Salud Carlos III (ISCIII) through project FIS-PI21/01642, co-funded by the European Union.

## AUTHOR’S CONTRIBUTIONS

Conceptualisation: A.F.G.J., E.L.G., H.T.R. Experiments: A.F.G.J., A.P., A.S.C. Formal analysis: A.F.G.J., E.V. Material support: A.B., A.D., V.L., L.I.G.G. Supervision: O.C., E.L.G., H.T.R. Funding acquisition: H.T.R. Writing-review and editing: A.F.G.J., H.T.R.

## ACKNOWLEDGEMENTS

We thank patients and family members for kindly donating blood samples and Dr. Mar Valés-Gómez (CNB-CSIC) for reagents and advice. Álvaro F. García-Jiménez was a graduate student in the Molecular Biosciences doctoral program of the Autonomous University of Madrid.

## SUPPLEMENTARY LEGENDS

Supplementary Table 1. Cell count per cluster, status, and sample within the proliferation-based scRNA-seq.

Supplementary Table 2. Cell count per cluster, status, and sample within the activation-based scRNA-seq.

Supplementary Table 3. Flow cytometry reagents.

Supplementary Figure 1. *In vitro* assays proving EBV stimulation of immune cells. (A) FACS dot-plots of IFN-γ and TNF-α producing T cells in XMEN or (B) XIAP deficient samples after incubation with EBV for 7 days, rest with IL-2 for 5 days, and restimulation with EBV or HCMV peptides at day 12. (C) Dot-plot showing the percentage of CD4, CD8, NK, and NKT cells within CTLs from healthy donors. (D) CD4 and CD8 staining of XMEN-derived CTLs. (E) Upregulation of LAMP1 in XMEN CTLs after incubation with autologous LCLs or PMA/ionomycin. (F) Specific-killing assay between XMEN CTLs and autologous LCLs.

Supplementary Figure 2. Characteristics of proliferating cells captured in scRNA-seq (I). (A) Cell sorting of PI^-^CTV^low^ cells. (B) Production of IFN-γ and TNF-α in the “EBV- naive” sample after stimulation for 6 days and restimulation for 4 hours with EBV or HCMV peptides. (C) UMAP distribution of cells split per status. (D) UMAP distribution of cells with detection of EBV transcripts split per status. The percentage of infected cells within the dataset is highlighted in red.

Supplementary Figure 3. Characteristics of proliferating cells captured in scRNA-seq (I). (A) Stacked bar-plot showing the distribution of CD4 clusters per status. (B) Stacked bar-plots depicting clonal expansion within CD4 clusters per status. (C) Representative examples of expanded CD4 clones per status. (D) Feature plot coloured by *FCGR3A* expression within the C9 UMAP. (E) UMAP coloured by clonal expansion within the C9 UMAP. (F) Representative examples of expanded CD8 clones per status. (G) Stacked bar-plot showing the distribution of CD8 subclusters per status. (H) Matrix plot showing Pearson’s correlation between the subclusters of CD8. (I) Violin-plot featuring the expression of *MKI67* within CD8 subclusters. (J) Violin-plots featuring the expression of CX3CR1, FGFBP2, FCGR3A, NCAM1, COTL1, and XCL1 genes within NK clusters. (K) Feature-plots showing the double expression of *TRDV2*/*TRGV9* and *TRAV1-2*/*KLRB1* within C10.

Supplementary Figure 4. Characteristics of activated cells against EBV (I). (A) Cell sorting of PI^-^ PBMCs expressing CD69, CD25, or CD69+CD25 in similar proportions. (B) Detection of activation markers (CD69 and/or CD25) in CD4, CD8, and NK cell populations at days 3, 4, 5, or 6 after incubation with EBV. (C) Detection of PD-1 within CD25^+^ or CD25^-^ fractions of CD4, CD8, or NK cells at day 3, 4, 5, or 6 after incubation with EBV. (D) Pseudobulk PCA of samples sequenced in the activation-based scRNA-seq coloured by condition. (E) Heatmap showing the bulk expression of the top4 markers of each cluster according to the Z-score calculated on the variable features of the whole dataset.

Supplementary Figure 5. Characteristics of activated cells against EBV (II). (A) Stacked bar-plot showing the distribution of CD4 T cell subclusters per mutation. (B) Heatmap showing the top5 differentially expressed genes per CD4 T cell subcluster. (C) Distribution of clonotype abundances per mutation within CD4 T cells. (D) Heatmap showing the top5 differentially expressed genes per CD8 T cell subcluster. (E) Violin-plots featuring the expression of *ZNF683*, *FGFBP2*, *GZMK*, *XCL1*, *GNLY*, *GZMB*, *KLRD1*, *KLRC2*, *KLRF1*, *KLRC3*, *LILRB1*, and *CD160* genes within CD8 T cell SC5-SC10. (F) UMAP coloured by individual clone size within CD8 T cells. (G) Heatmaps of *TRAV* and *TRBV* gene usage per mutation within CD8 T cells. (H) UMAP of CD8 T cells coloured by EBV antigen TCRβ specificity according to VDJdb.

Supplementary Figure 6. Characteristics of activated cells against EBV (III). (A) UMAP distribution of NK cells at 0.5 resolution. (B) Stacked bar-plot showing the distribution of NK cell subclusters per mutation. (C) Feature plots coloured by *NCAM1* and *FGFBP2* expression, highlighting regions that define CD56^bright^ and CD56^dim^ cells, respectively. (D) Violin plots showing notable differences between HD and XMEN in CD56^dim^ cells. (E) Stacked violin plot of specific markers upregulated or downregulated in adaptive NK cells.

Supplementary Figure 7. Characteristics of activated cells against EBV (IV). (A) UMAP distribution of Vδ2 cells at 0.5 resolution and UMAP grouped by mutation. (B) Heatmap showing the top5 differential expressed genes per Vδ2 subcluster. (C) UMAP distribution of unconventional Vδ cells at 0.5 resolution split by mutation. (D) Heatmap showing the top5 differential expressed genes per unconventional Vδ subcluster. (E) Violin-plots featuring the expression of CX3CR1, KLRG1, and GZMK within uncoventional Vδ subclusters. (F) Feature plots coloured by *GNLY* and *GZMB* expression for healthy and XMEN uncoventional Vδ cells. (G) UMAP distribution of MAIT cells at 0.5 resolution. (H) UMAP distribution of MAIT cells grouped by mutation. (I) Stacked bar-plots showing VJ and VDJ usage of MAIT cells. (J) Stacked barplots depicting clonal expansion within MAIT subclusters. (K) Violin-plots of MAIT activation markers according to De Biasi *et al*^27^.

Supplementary Figure 8. *In vitro* characterisation of cell populations (I). (A) Monitoring of a rapid decrease in the B cell population at day 1, 2, or 3 after incubation with EBV. (B) Expression of GM-CSF within CCR2-expressing CD4 T cells after stimulation with EBV at day 3. (C) T cell panel of antibodies and gating of input cells (CD3^+^). (D) Upregulation of CCR5 within CD4 and CD8 T cells after stimulation of PBMCs with EBV + IL-2/12/15. (E) CD8 expression in Vδ1 cells. (F) NKG2C expression in CD69- expressing Vδ1 cells. (G) Upregulation of KIR2DL3 and CD16 after incubation with EBV. (H) Expansion of Vδ2 cells after treating PBMCs with EBV + IL-2/12/15. (I) Representative example of the gating of MAIT cells (CD3^+^CD161^++^TCRαv7.2^+^) and the dilution of CTV in control and experimental conditions. Bar-plot showing mean value ± SEM. (J) Vδ1, Vδ2, and MAIT panel of antibodies and gating of input cells (CD3^+^TRDV1^+^, CD3^+^TRDV2^+^ and CD3^+^CD161^+^TCRVα7.2^+^). (K) NK cell panel of antibodies and gating of input cells (CD3^-^CD56^+^).

Supplementary Figure 9. Features of *in vitro* expanded Vδ1 cells. (A) Example of the natural upregulation of TRDV1 molecule after 6 days of EBV incubation of PBMCs. (B) Stacked violin plots representing the expression of different markers in Vδ1 cells from SC1 and SC2. (C) Example of expansion of DOT cells following a 3-4 week protocol. (D) Examples of LAMP1 signals after incubation with LCLs in non-resting and resting conditions. (E) Specific killing of LCLs in resting condition when using expanded DOT cells at a 5:1 and 10:1 ratio, measured through the release of calcein molecule. (F) Example of LAMP1 degranulation after using blocking antibodies in DOT cells against NKG2D, NKp30, NKG2C, NKp44, and/or DNAM-1. (G) Detection of LAMP1 after blocking TRDV1. (H) Bar-plot and histograms representing the MFI of B7-H6, HLA-E, MICA, MICB, ULBP1, ULBP2, CD155, or Nectin-2 within LCLs. (I) Histograms of B7-H6 and MICA expresion in Mutu, Kem, and Raji cell lines. (J) Example of LAMP1 degranulation after using blocking antibodies in LCLs against B7-H6, MICA, ULBP1, and/or CD155. (K) Histograms representing the detection of CD27, NKG2D, NKp30, NKG2C, NKp44, and DNAM-1 in expanded DOT cells in the resting, resting + IL-15, and non-resting conditions.

Supplementary Figure 10. DOT cells *in vivo*. (A) Kinetics of the expansion of Delta One T (DOT) cells. (B) Percentage of TRDV1^+^ events and detection of CD27, NKp30, and NKG2D for the donors used in the *in vivo* experiment. (C) Graph representing the weight of mice through the experiment in tumour-only (n=3) and tumour + DOT (n=3) conditions. (D) Representative flow cytometry plots and quantification of DOT cells expressing CD62L and/or CD45RA in blood, spleen, and tumour at sacrifice. Paired sample t-test (*p<0.05) (E) Tumour volume fold change (FC) in tumour-only (n=3) and tumour + DOT (n=3) conditions. The volume on day 4 was used as reference with a FC value of 1.

Supplementary Figure 11. Additional characterisation of samples. (A) Pedigree of the 3 samples with mutations in *MAGT1*. (B) Determination of the inverted CD4/CD8 ratio in XMEN samples. (C) Detection of NKG2D in the CD8 T cells of healthy and XMEN samples. (D) Paired-plots representing different conditions for EBV-stimulation of PBMCs. Paired-sample t-test (*p<0.05, **p<0.01, ***p<0.001).

## Notes

https://www.ncbi.nlm.nih.gov/geo/query/acc.cgi?acc=GSE277638

https://github.com/algarji/PBMCs_EBV_scRNA-seq.

### Competing Interest Statement

The authors have declared no competing interest.

