## Supplementary tables and data for "A high-resolution, unbiased analysis of the cellular immune response to Epstein-Barr virus"

### Supp. Table 1

| | B518 | B618 | B204 | B108 | XMEN_1 | XMEN_2 | XMEN_3 | XIAP_1 | Total |
|---|---|---|---|---|---|---|---|---|---|
| | Healthy Donors | | | EBV_naive | MAGT1$^{y/-}$ | | | XIAP$^{y/-}$ | |
| C1 | 795 | 42 | 229 | 21 | 20 | 13 | 1 | 1 | 1122 |
| C2 | 35 | 62 | 19 | 118 | 352 | 75 | 1 | 111 | 773 |
| C3 | 35 | 4 | 360 | 3 | 17 | 0 | 0 | 61 | 480 |
| C4 | 38 | 45 | 39 | 126 | 170 | 48 | 0 | 241 | 707 |
| C5 | 380 | 96 | 2 | 5 | 3 | 1 | 0 | 0 | 487 |
| C6 | 12 | 58 | 35 | 337 | 95 | 11 | 5 | 5 | 558 |
| C7 | 29 | 6 | 153 | 7 | 66 | 7 | 0 | 143 | 411 |
| C8 | 11 | 56 | 7 | 290 | 2 | 1 | 0 | 3 | 370 |
| C9 | 91 | 61 | 30 | 7 | 104 | 31 | 6 | 1 | 331 |
| C10 | 184 | 10 | 21 | 3 | 22 | 3 | 2 | 0 | 245 |
| C11 | 74 | 25 | 10 | 85 | 4 | 1 | 0 | 60 | 259 |
| C12 | 173 | 9 | 57 | 4 | 3 | 9 | 1 | 2 | 258 |
| C13 | 76 | 39 | 9 | 103 | 0 | 0 | 0 | 22 | 249 |
| C14 | 40 | 13 | 9 | 56 | 5 | 3 | 0 | 27 | 153 |
| C15 | 21 | 0 | 114 | 0 | 22 | 0 | 1 | 32 | 190 |
| C16 | 3 | 11 | 15 | 13 | 38 | 15 | 1 | 0 | 96 |
| C17 | 14 | 4 | 10 | 2 | 18 | 1 | 2 | 1 | 52 |
| C18 | 29 | 8 | 46 | 2 | 14 | 0 | 0 | 23 | 122 |
| Total | 2040 | 549 | 1165 | 1182 | 955 | 219 | 20 | 733 | 6863 |

### Supp. Table 2

| | HD080 | HD1268 | HD756 | HD04 | HD05 | HD06 | HD07 | HD08 | HD09 | XMEN_1 | XMEN_3 | XIAP_2 | PIK3CD_1 | PIK3CD_2 |
|---|---|---|---|---|---|---|---|---|---|---|---|---|---|---|
| | Healthy donors | | | | | | | | | XMEN | | | APDS | |
| C1 | 157 | 84 | 99 | 86 | 96 | 47 | 99 | 65 | 44 | 309 | 2599 | 231 | 1871 | 296 |
| C2 | 241 | 332 | 171 | 144 | 280 | 102 | 162 | 34 | 65 | 330 | 1143 | 5 | 148 | 1136 |
| C3 | 248 | 69 | 344 | 765 | 180 | 545 | 82 | 58 | 42 | 86 | 638 | 751 | 274 | 588 |
| C4 | 85 | 65 | 157 | 572 | 25 | 86 | 45 | 31 | 13 | 61 | 261 | 1639 | 123 | 82 |
| C5 | 549 | 608 | 764 | 117 | 59 | 24 | 59 | 133 | 39 | 98 | 289 | 18 | 122 | 59 |
| C6 | 113 | 86 | 154 | 49 | 47 | 62 | 133 | 37 | 46 | 264 | 649 | 215 | 301 | 164 |
| C7 | 178 | 90 | 174 | 630 | 71 | 102 | 45 | 23 | 24 | 61 | 129 | 204 | 40 | 46 |
| C8 | 104 | 114 | 78 | 47 | 19 | 15 | 158 | 335 | 115 | 39 | 132 | 43 | 53 | 40 |
| C9 | 290 | 105 | 217 | 136 | 77 | 69 | 40 | 15 | 31 | 35 | 94 | 19 | 375 | 129 |
| C10 | 110 | 289 | 19 | 49 | 92 | 56 | 23 | 24 | 16 | 231 | 235 | 3 | 15 | 223 |
| C11 | 12 | 44 | 142 | 45 | 54 | 15 | 21 | 26 | 37 | 205 | 325 | 6 | 86 | 57 |
| C12 | 39 | 7 | 497 | 172 | 85 | 115 | 37 | 97 | 46 | 1 | 5 | 0 | 0 | 0 |
| C13 | 33 | 4 | 281 | 23 | 3 | 2 | 24 | 20 | 47 | 2 | 0 | 0 | 0 | 0 |
| C14 | 2 | 0 | 0 | 4 | 4 | 1 | 1 | 2 | 0 | 31 | 310 | 20 | 127 | 4 |
| C15 | 36 | 17 | 24 | 20 | 14 | 4 | 9 | 19 | 4 | 36 | 93 | 2 | 7 | 8 |
| C16 | 6 | 75 | 6 | 8 | 14 | 4 | 5 | 3 | 2 | 33 | 112 | 21 | 12 | 4 |
| C17 | 19 | 73 | 22 | 4 | 3 | 1 | 2 | 0 | 2 | 13 | 23 | 0 | 1 | 0 |
| C18 | 0 | 0 | 1 | 0 | 0 | 0 | 8 | 12 | 17 | 0 | 0 | 0 | 0 | 0 |
| C19 | 2 | 1 | 3 | 17 | 3 | 8 | 3 | 1 | 1 | 3 | 13 | 22 | 3 | 5 |
| C20 | 1 | 12 | 56 | 2 | 0 | 0 | 1 | 0 | 0 | 1 | 1 | 3 | 4 | 1 |
| C21 | 16 | 1 | 10 | 0 | 2 | 3 | 0 | 0 | 2 | 5 | 5 | 0 | 4 | 2 |

### Supp. Table 3

| | Fluorophore | Clone | Supplier |
|---|---|---|---|
| Viability | | | |
| AnnexinV | APC | - | BD Biosciences |
| AnnexinV | PE | - | BD Biosciences |
| DAPI | - | - | BD Biosciences |
| LIVE/DEAD Fixable Blue | - | - | Thermo Fisher |
| Propidium iodide | - | - | BioLegend |
| Zombie Aqua Fixable | - | - | BioLegend |
| Surface | | | |
| CCR2 | APC | K036C2 | BioLegend |
| CCR5 | PB450 | HEK/1/85a | BioLegend |
| CD16 | APC | 3G8 | BioLegend |
| CD16 | V500 | 3G8 | BD Horizon |
| CD161 | PE | W18070C | BioLegend |
| CD161 | AF700 | HP-3G10 | BioLegend |
| CD19 | BYG667 | SJ25C1 | CYTEK |
| CD19 | FITC | HIB19 | BioLegend |
| CD25 | PB450 | BC96 | BioLegend |
| CD25 | APC-Fire810 | M-A251 | BioLegend |
| CD25 | PE | BC96 | BioLegend |
| CD25 | PE/Cy7 | BC96 | BioLegend |
| CD27 | FITC | M-T271 | BioLegend |
| CD3 | BUV496 | UCHT1 | BD Biosciences |
| CD3 | FITC | UCHT1 | BioLegend |
| CD3 | PB450 | UCHT1 | BioLegend |
| CD3 | PerCP5.5 | OKT3 | BioLegend |
| CD38 | Spark-NIR685 | HIT2 | BioLegend |
| CD39 | FITC | A1 | BioLegend |
| CD4 | APC | OKT4 | BioLegend |
| CD4 | FITC | OKT4 | BioLegend |
| CD4 | V500 | RPA-T4 | BD Biosciences |
| CD45 | BUV395 | HI30 | BD Biosciences |
| CD45RA | PB450 | HI100 | BioLegend |
| CD56 | PE | MEM-188 | BioLegend |
| CD56 | PE-Cy5 | MEM-188 | BioLegend |
| CD56 | PE-Dazzle594 | 5.1H11 | BioLegend |
| CD57 | PE-Dazzle594 | HNK-1 | BioLegend |
| CD62L | PE-Cy7 | DREG-56 | BioLegend |
| CD69 | Alexa Fluor 700 | FN50 | BioLegend |
| CD69 | APC/Cy7 | FN50 | BioLegend |
| CD69 | PE/Cy7 | FN50 | BioLegend |
| CD8 | APC/Cy7 | SK1 | BioLegend |
| CD8 | PE/Cy7 | SK1 | BioLegend |
| CD94 | FITC | DX22 | BioLegend |
| CTV | - | - | ThermoFisher |
| CXCR3 | APC | G025H7 | BioLegend |
| HLA-DR | APC-Cy7 | L243 | BioLegend |
| HLA-DR | PE | LN3 | BioLegend |
| KIR2DL2/L3 | PE/Cy7 | DX27 | BioLegend |
| KIR2DL1/S1/S3/S5 | PE/Cy7 | HP-MA4 | BioLegend |
| KLRG1 | APC | SA231A2 | BioLegend |
| LAMP1 | APC | H4A3 | BioLegend |
| LAMP1 | PerCP | H4A3 | BioLegend |
| NKG2A | APC | S19004C | BioLegend |
| NKG2A | PE-Cy5 | S19004C | BioLegend |
| NKG2C | Alexa Fluor 488 | - | R&D Systems |
| NKG2D | PE | 1D11 | BioLegend |
| NKp30 | BV421 | P30-15 | BioLegend |
| NKp30 | APC | P30-15 | BioLegend |
| NKp44 | PE/Cy7 | P44-8 | BioLegend |
| NKp80 | APC | 5D12 | BioLegend |
| PD-1 | BUV737 | EH12.1 | BD Biosciences |
| PD-1 | APC/Cy7 | NAT105 | BioLegend |
| TCR Vα7.2 | FITC | 3C10 | BioLegend |
| TCRVα7.2 | PE-Cy7 | 3C10 | BioLegend |
| TRDV1 | PE | TS8.2 | ThermoFisher |
| TRDV2 | APC | B6 | BioLegend |
| Intracellular | | | |
| GM-CSF | PE | BVD2-21C11 | BioLegend |
| IFNγ | APC | 4S.B3 | BioLegend |
| IFNγ | PE/Cy7 | 4S.B3 | BioLegend |
| GZMB | PE-Dazzle594 | QA18A28 | BioLegend |
| GZMB | PerCP5.5 | QA16A02 | BioLegend |
| TNFα | APC | MAb11 | BioLegend |
| TNFα | PE | MAb11 | BioLegend |

**Supp. 1**

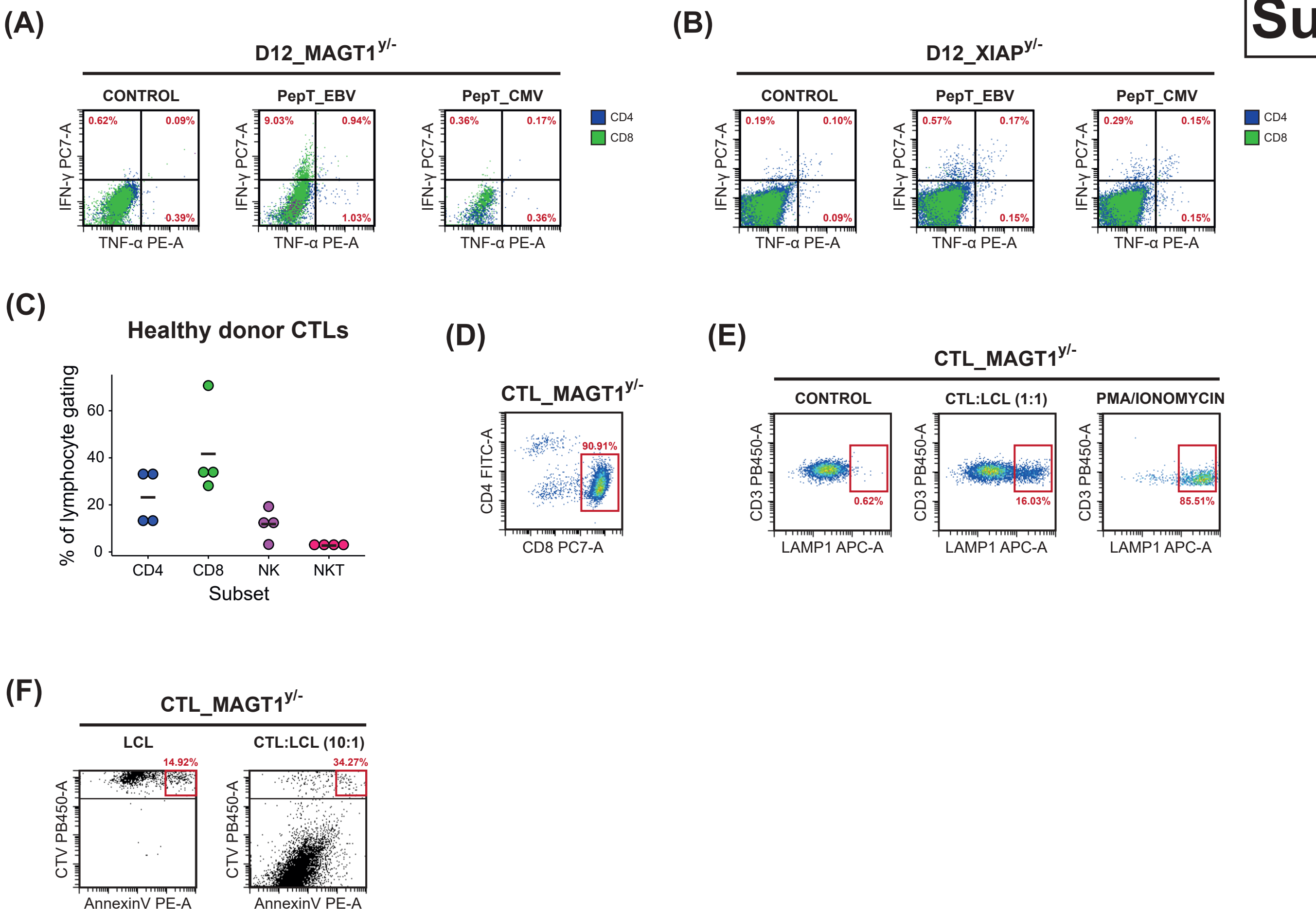

(A)
D12_MAGT1y/-
CONTROL
PepT_EBV
PepT_CMV
0.62%
0.09%
0.39%
9.03%
0.94%
1.03%
0.36%
0.17%
0.36%
IFN-γ PC7-A
TNF-α PE-A
CD4
CD8
(B)
D12_XIAPy/-
CONTROL
PepT_EBV
PepT_CMV
0.19%
0.10%
0.09%
0.57%
0.17%
0.15%
0.29%
0.15%
0.15%
IFN-γ PC7-A
TNF-α PE-A
CD4
CD8
(C)
Healthy donor CTLs
% of lymphocyte gating
0
20
40
60
CD4
CD8
NK
NKT
Subset
(D)
CTL_MAGT1y/-
CD4 FITC-A
CD8 PC7-A
90.91%
(E)
CTL_MAGT1y/-
CONTROL
CTL:LCL (1:1)
PMA/IONOMYCIN
CD3 PB450-A
LAMP1 APC-A
0.62%
16.03%
85.51%
(F)
CTL_MAGT1y/-
LCL
CTL:LCL (10:1)
14.92%
34.27%
CTV PB450-A
AnnexinV PE-A

**Supp. 2**

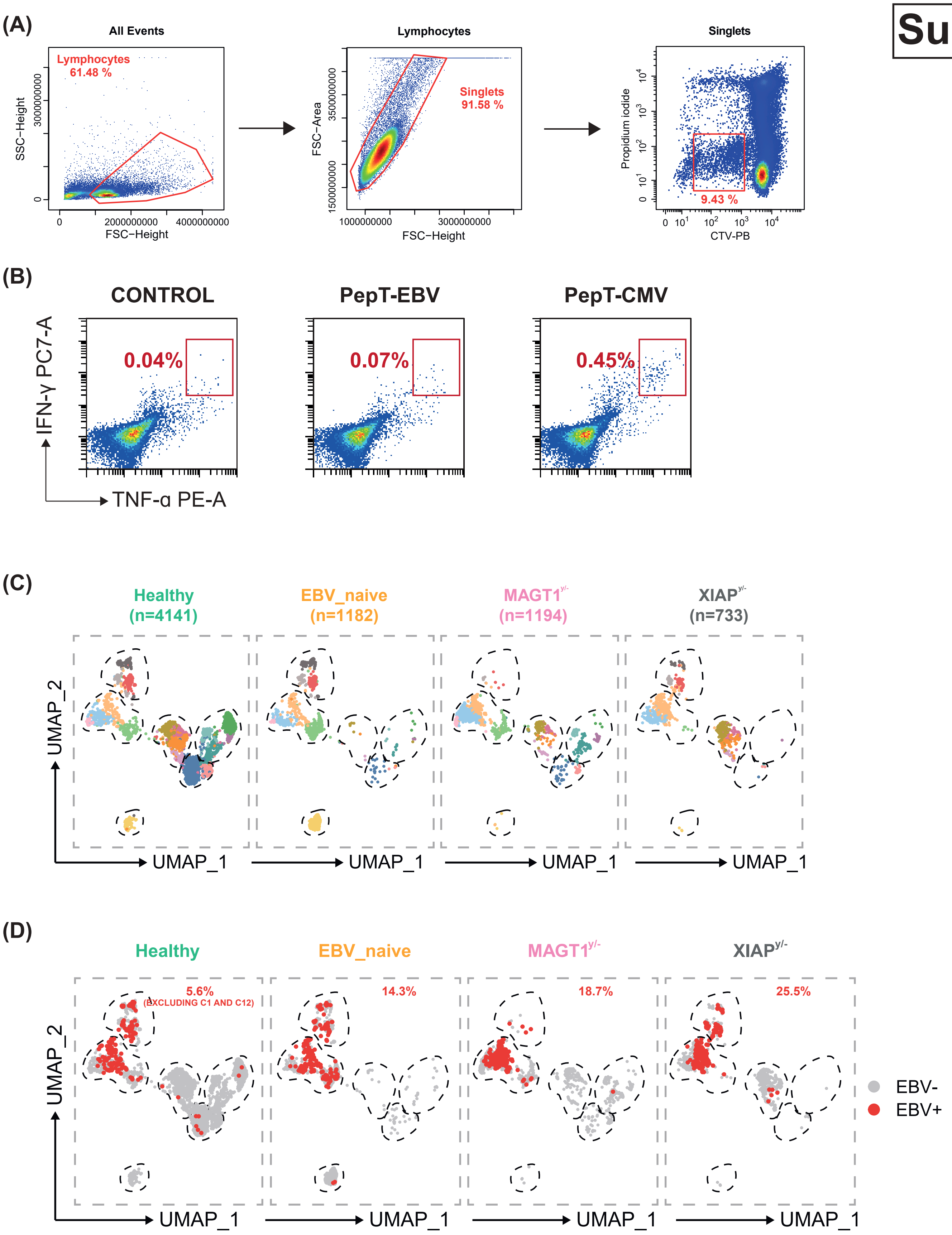
(A)
All Events
Lymphocytes
61.48 %
SSC−Height
FSC−Height
Lymphocytes
Singlets
91.58 %
FSC−Area
FSC−Height
Singlets
Propidium iodide
9.43 %
CTV-PB
(B)
CONTROL
PepT-EBV
PepT-CMV
0.04%
0.07%
0.45%
IFN-γ PC7-A
TNF-α PE-A
(C)
Healthy (n=4141)
EBV_naive (n=1182)
MAGT1y/- (n=1194)
XIAPy/- (n=733)
UMAP_2
UMAP_1
(D)
Healthy
EBV_naive
MAGT1y/-
XIAPy/-
5.6%
(EXCLUDING C1 AND C12)
14.3%
18.7%
25.5%
EBV-
EBV+
UMAP_2
UMAP_1

Supp. 3

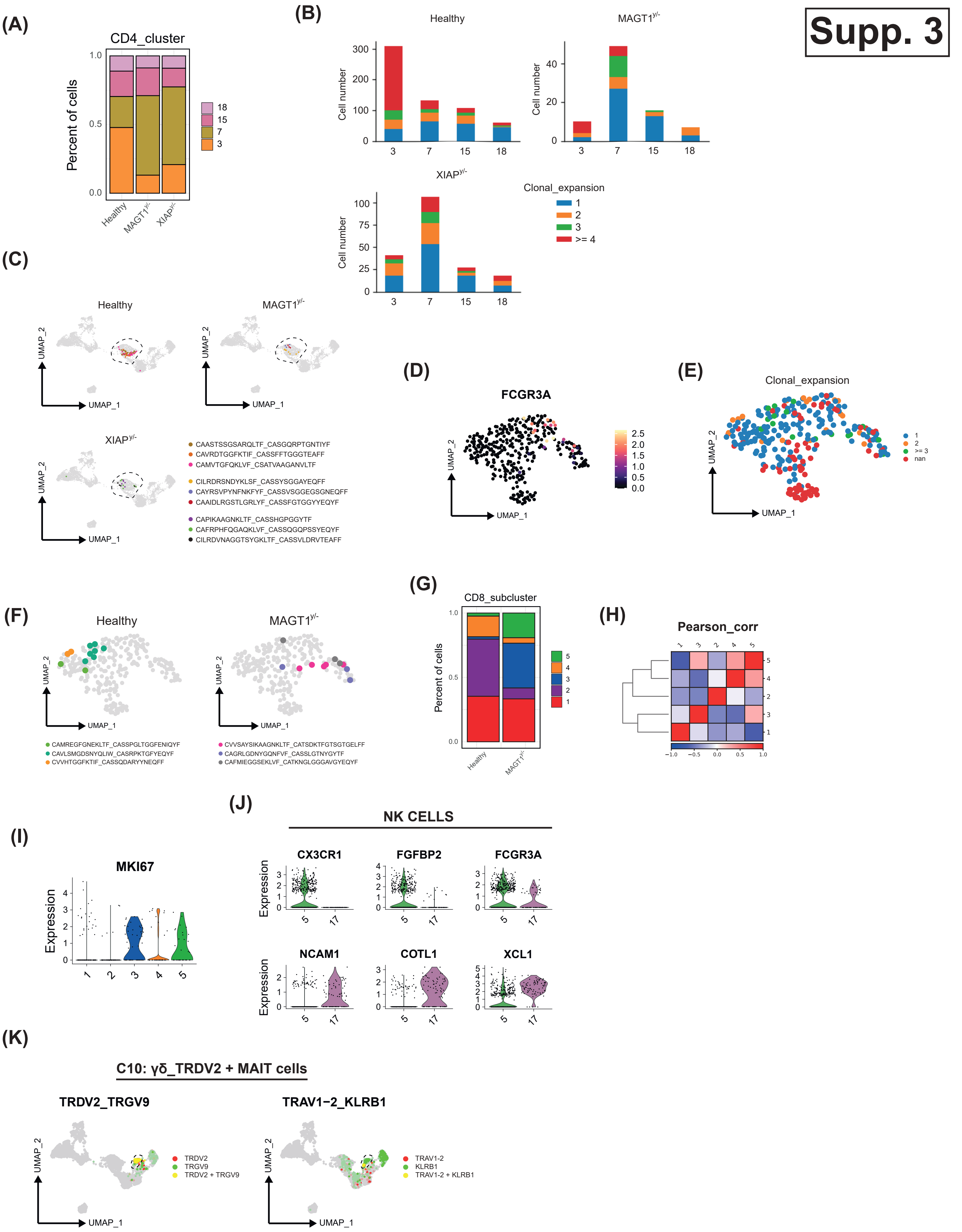
(A)
CD4_cluster
Percent of cells
1.0
0.5
0.0
Healthy
MAGT1y/-
XIAPy/-
18
15
7
3
(B)
Healthy
MAGT1y/-
XIAPy/-
Cell number
3
7
15
18
Clonal_expansion
1
2
3
>= 4
(C)
Healthy
MAGT1y/-
XIAPy/-
UMAP_1
UMAP_2
CAASTSSGSARQLTF_CASGQRPTGNTIYF
CAVRDTGGFKTIF_CASSFFTGGGTEAFF
CAMVTGFQKLVF_CSATVAAGANVLTF
CILRDRSNDYKLSF_CASSYSGGAYEQFF
CAYRSVPYNFNKFYF_CASSVSGGEGSGNEQFF
CAAIDLRGSTLGRLYF_CASSFGTGGYYEQYF
CAPIKAAGNKLTF_CASSHGPGGYTF
CAFRPHFQGAQKLVF_CASSQGQPSSYEQYF
CILRDVNAGGTSYGKLTF_CASSVLDRVTEAFF
(D)
FCGR3A
UMAP_1
UMAP_2
2.5
2.0
1.5
1.0
0.5
0.0
(E)
Clonal_expansion
UMAP_1
UMAP_2
1
2
>= 3
nan
(F)
Healthy
MAGT1y/-
UMAP_1
UMAP_2
CAMREGFGNEKLTF_CASSPGLTGGFENIQYF
CAVLSMGDSNYQLIW_CASRPKTGFYEQYF
CVVHTGGFKTIF_CASSQDARYYNEQFF
CVVSAYSIKAAGNKLTF_CATSDKTFGTSGTGELFF
CAGRLGDNYGQNFVF_CASSLGTNYGYTF
CAFMIEGGSEKLVF_CATKNGLGGGAVGYEQYF
(G)
CD8_subcluster
Percent of cells
1.0
0.5
0.0
Healthy
MAGT1y/-
5
4
3
2
1
(H)
Pearson_corr
1
3
2
4
5
−1.0
−0.5
0.0
0.5
1.0
(I)
MKI67
Expression
1
2
3
4
5
(J)
NK CELLS
CX3CR1
FGFBP2
FCGR3A
NCAM1
COTL1
XCL1
Expression
5
17
(K)
C10: γδ_TRDV2 + MAIT cells
TRDV2_TRGV9
TRAV1−2_KLRB1
UMAP_1
UMAP_2
TRDV2
TRGV9
TRDV2 + TRGV9
TRAV1-2
KLRB1
TRAV1-2 + KLRB1

Supp. 4

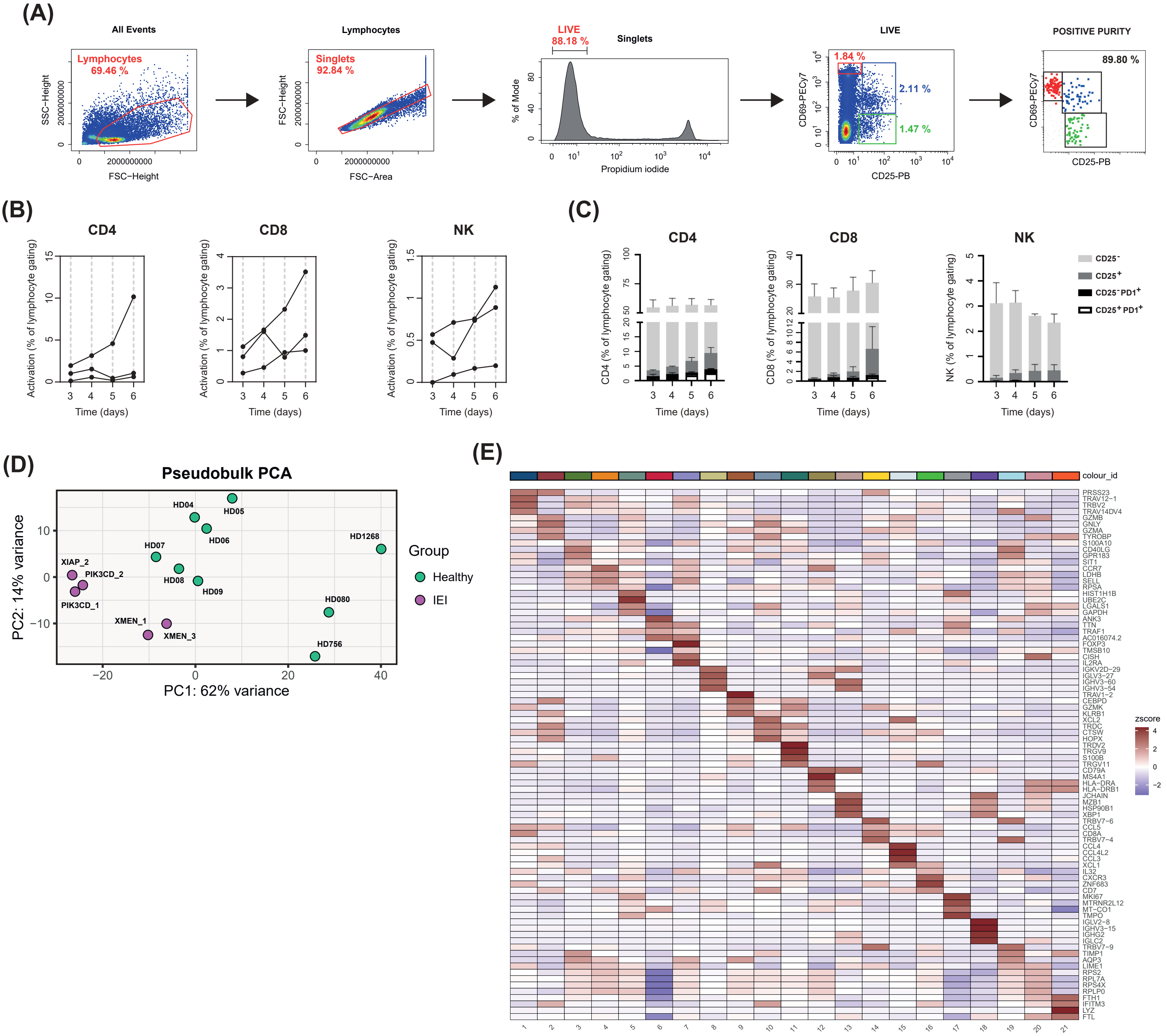
(A)
All Events
Lymphocytes 69.46 %
SSC−Height
FSC−Height
Lymphocytes
Singlets 92.84 %
FSC−Height
FSC−Area
LIVE 88.18 %
Singlets
% of Mode
Propidium iodide
LIVE
1.84 %
2.11 %
1.47 %
CD69-PECy7
CD25-PB
POSITIVE PURITY
89.80 %
CD69-PECy7
CD25-PB
(B)
CD4
CD8
NK
Activation (% of lymphocyte gating)
Time (days)
(C)
CD4
CD8
NK
CD4 (% of lymphocyte gating)
CD8 (% of lymphocyte gating)
NK (% of lymphocyte gating)
Time (days)
CD25⁻
CD25⁺
CD25⁻PD1⁺
CD25⁺PD1⁺
(D)
Pseudobulk PCA
PC2: 14% variance
PC1: 62% variance
HD04
HD05
HD06
HD07
HD08
HD09
HD1268
HD080
HD756
XIAP_2
PIK3CD_2
PIK3CD_1
XMEN_1
XMEN_3
Group
Healthy
IEI
(E)
colour_id
zscore
PRSS23
TRAV12−1
TRBV2
TRAV14DV4
GZMB
GNLY
GZMA
TYROBP
S100A10
CD40LG
GPR183
SIT1
CCR7
LDHB
SELL
RPSA
HIST1H1B
UBE2C
LGALS1
GAPDH
ANK3
TTN
TRAF1
AC016074.2
FOXP3
TMSB10
CISH
IL2RA
IGKV2D−29
IGLV3−27
IGHV3−60
IGHV3−54
TRAV1−2
CEBPD
GZMK
KLRB1
XCL2
TRDC
CTSW
HOPX
TRDV2
TRGV9
S100B
TRGV11
CD79A
MS4A1
HLA−DRA
HLA−DRB1
JCHAIN
MZB1
HSP90B1
XBP1
TRBV7−6
CCL5
CD8A
TRBV7−4
CCL4
CCL4L2
CCL3
XCL1
IL32
CXCR3
ZNF683
CD7
MKI67
MTRNR2L12
MT−CO1
TMPO
IGLV2−8
IGHV3−15
IGHG2
IGLC2
TRBV7−9
TIMP1
AQP3
LIME1
RPS2
RPL7A
RPS4X
RPLP0
FTH1
IFITM3
LYZ
FTL

**Supp. 5**

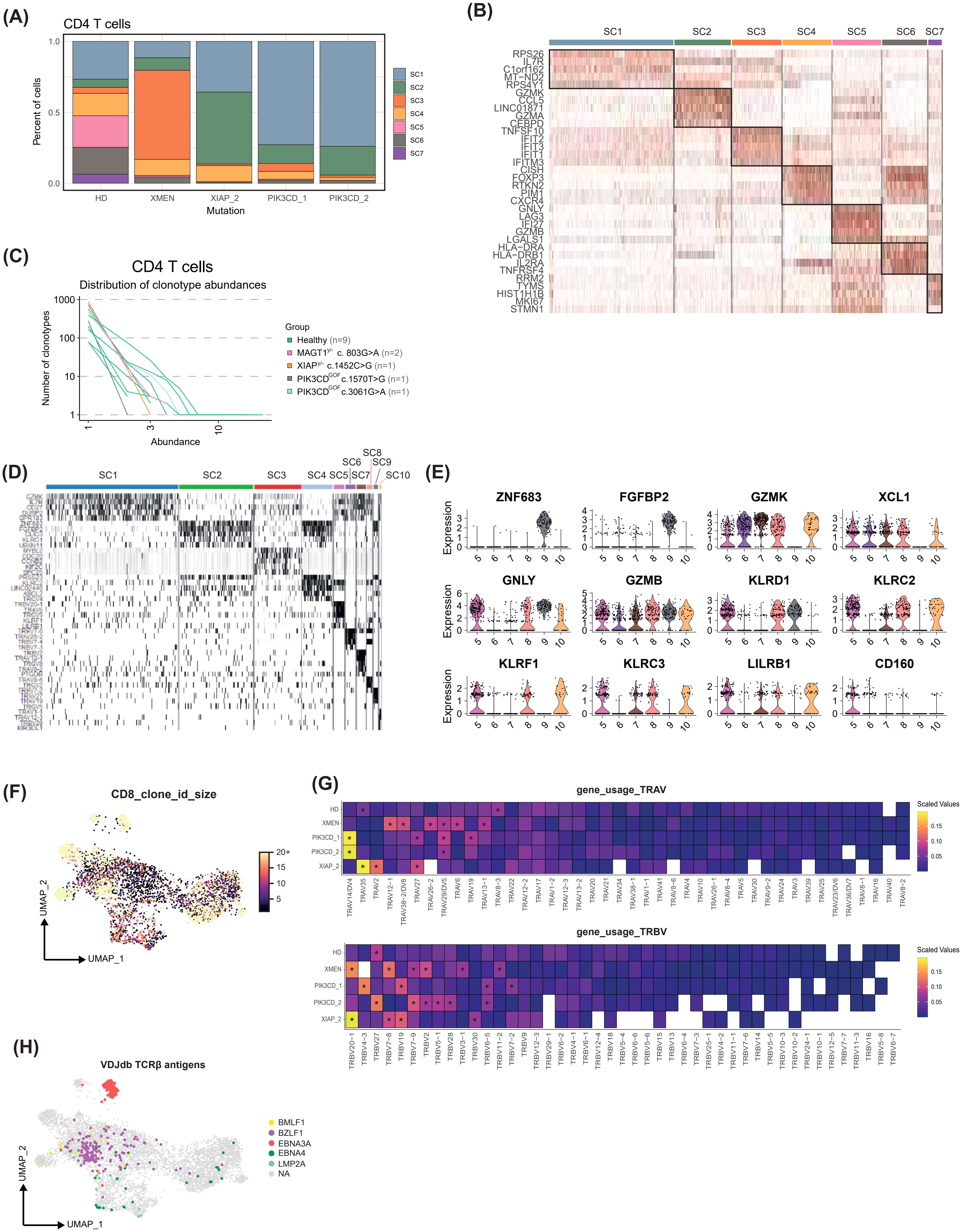
(A)
CD4 T cells
Percent of cells
1.0
0.5
0.0
HD
XMEN
XIAP_2
PIK3CD_1
PIK3CD_2
Mutation
SC1
SC2
SC3
SC4
SC5
SC6
SC7
(B)
SC1
SC2
SC3
SC4
SC5
SC6
SC7
RPS26
IL7R
C1orf162
MT-ND2
RPS4Y1
GZMK
CCL5
LINC01871
GZMA
CEBPD
TNFSF10
IFIT2
IFIT3
IFIT1
IFITM3
CISH
FOXP3
RTKN2
PIM1
CXCR4
GNLY
LAG3
IFI27
GZMB
LGALS1
HLA-DRA
HLA-DRB1
IL2RA
TNFRSF4
RRM2
TYMS
HIST1H1B
MKI67
STMN1
(C)
CD4 T cells
Distribution of clonotype abundances
Number of clonotypes
1000
100
10
1
1
3
10
Abundance
Group
Healthy (n=9)
MAGT1y/- c. 803G>A (n=2)
XIAPy/- c.1452C>G (n=1)
PIK3CDGOF c.1570T>G (n=1)
PIK3CDGOF c.3061G>A (n=1)
(D)
SC1
SC2
SC3
SC4
SC5
SC6
SC7
SC8
SC9
SC10
(E)
ZNF683
FGFBP2
GZMK
XCL1
GNLY
GZMB
KLRD1
KLRC2
KLRF1
KLRC3
LILRB1
CD160
Expression
5
6
7
8
9
10
(F)
CD8_clone_id_size
20+
15
10
5
UMAP_2
UMAP_1
(G)
gene_usage_TRAV
HD
XMEN
PIK3CD_1
PIK3CD_2
XIAP_2
TRAV14/DV4
TRAV35
TRAV2
TRAV12-1
TRAV38-2/DV8
TRAV27
TRAV26-2
TRAV29/DV5
TRAV6
TRAV19
TRAV13-1
TRAV8-3
TRAV22
TRAV12-2
TRAV17
TRAV1-2
TRAV12-3
TRAV13-2
TRAV20
TRAV21
TRAV34
TRAV38-1
TRAV1-1
TRAV41
TRAV8-6
TRAV4
TRAV10
TRAV26-1
TRAV8-4
TRAV5
TRAV30
TRAV9-2
TRAV24
TRAV3
TRAV39
TRAV25
TRAV23/DV6
TRAV36/DV7
TRAV8-1
TRAV16
TRAV40
TRAV8-2
Scaled Values
0.15
0.10
0.05
gene_usage_TRBV
HD
XMEN
PIK3CD_1
PIK3CD_2
XIAP_2
TRBV20-1
TRBV4-3
TRBV27
TRBV7-8
TRBV19
TRBV7-9
TRBV2
TRBV5-1
TRBV28
TRBV3-1
TRBV30
TRBV6-5
TRBV11-2
TRBV7-2
TRBV9
TRBV12-3
TRBV29-1
TRBV6-2
TRBV4-1
TRBV6-1
TRBV12-4
TRBV18
TRBV5-4
TRBV6-6
TRBV5-6
TRBV15
TRBV13
TRBV6-4
TRBV7-3
TRBV25-1
TRBV4-2
TRBV11-1
TRBV7-6
TRBV14
TRBV5-5
TRBV10-3
TRBV10-2
TRBV24-1
TRBV10-1
TRBV12-5
TRBV7-7
TRBV11-3
TRBV16
TRBV5-8
TRBV6-7
Scaled Values
0.15
0.10
0.05
(H)
VDJdb TCRβ antigens
BMLF1
BZLF1
EBNA3A
EBNA4
LMP2A
NA
UMAP_2
UMAP_1

**Supp. 6**

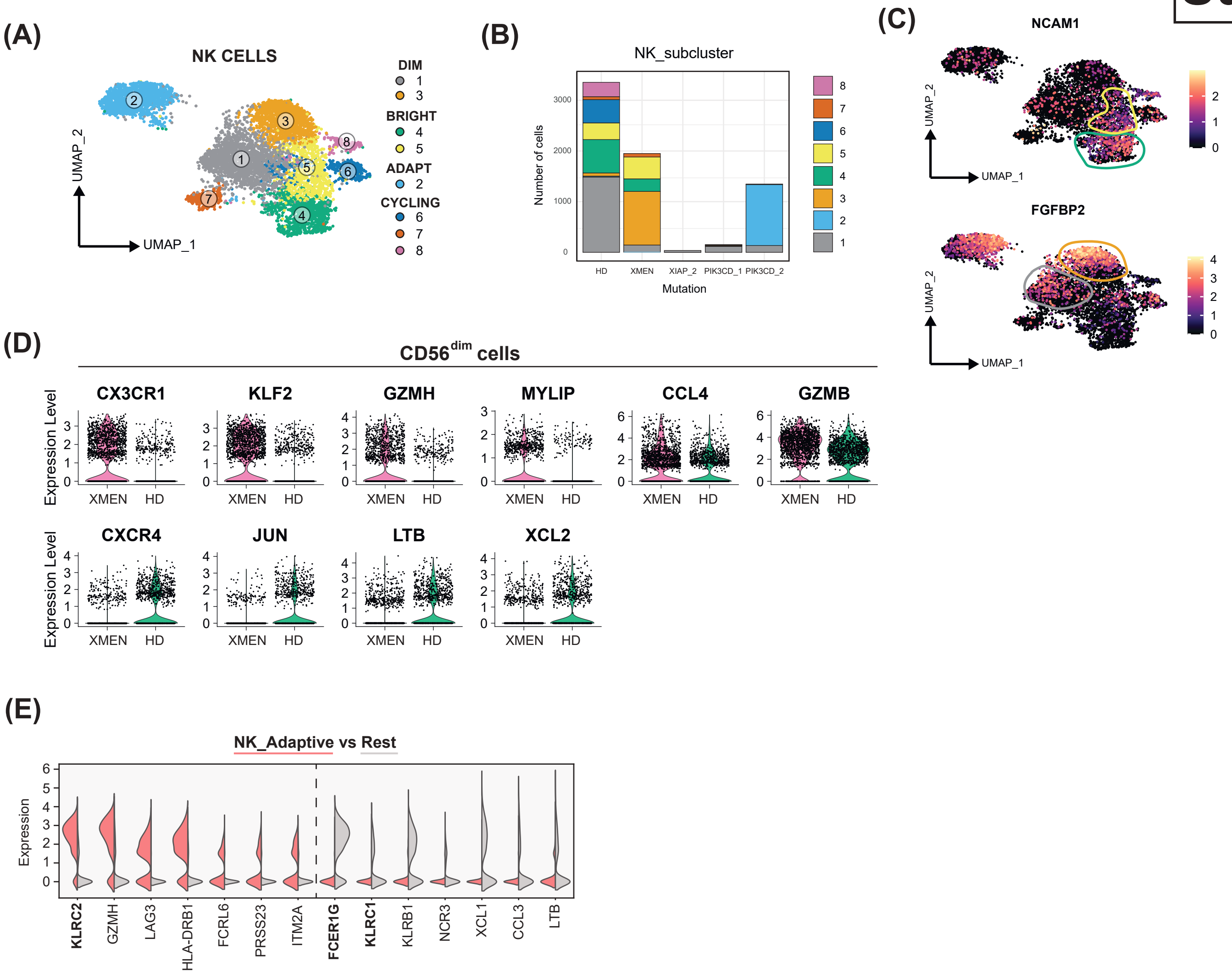
(A)
NK CELLS
UMAP_2
UMAP_1
DIM
1
3
BRIGHT
4
5
ADAPT
2
CYCLING
6
7
8
(B)
NK_subcluster
Number of cells
3000
2000
1000
0
HD
XMEN
XIAP_2
PIK3CD_1
PIK3CD_2
Mutation
8
7
6
5
4
3
2
1
(C)
NCAM1
FGFBP2
UMAP_2
UMAP_1
(D)
CD56dim cells
CX3CR1
KLF2
GZMH
MYLIP
CCL4
GZMB
CXCR4
JUN
LTB
XCL2
Expression Level
XMEN
HD
(E)
NK_Adaptive vs Rest
Expression
KLRC2
GZMH
LAG3
HLA-DRB1
FCRL6
PRSS23
ITM2A
FCER1G
KLRC1
KLRB1
NCR3
XCL1
CCL3
LTB

**Supp. 7**

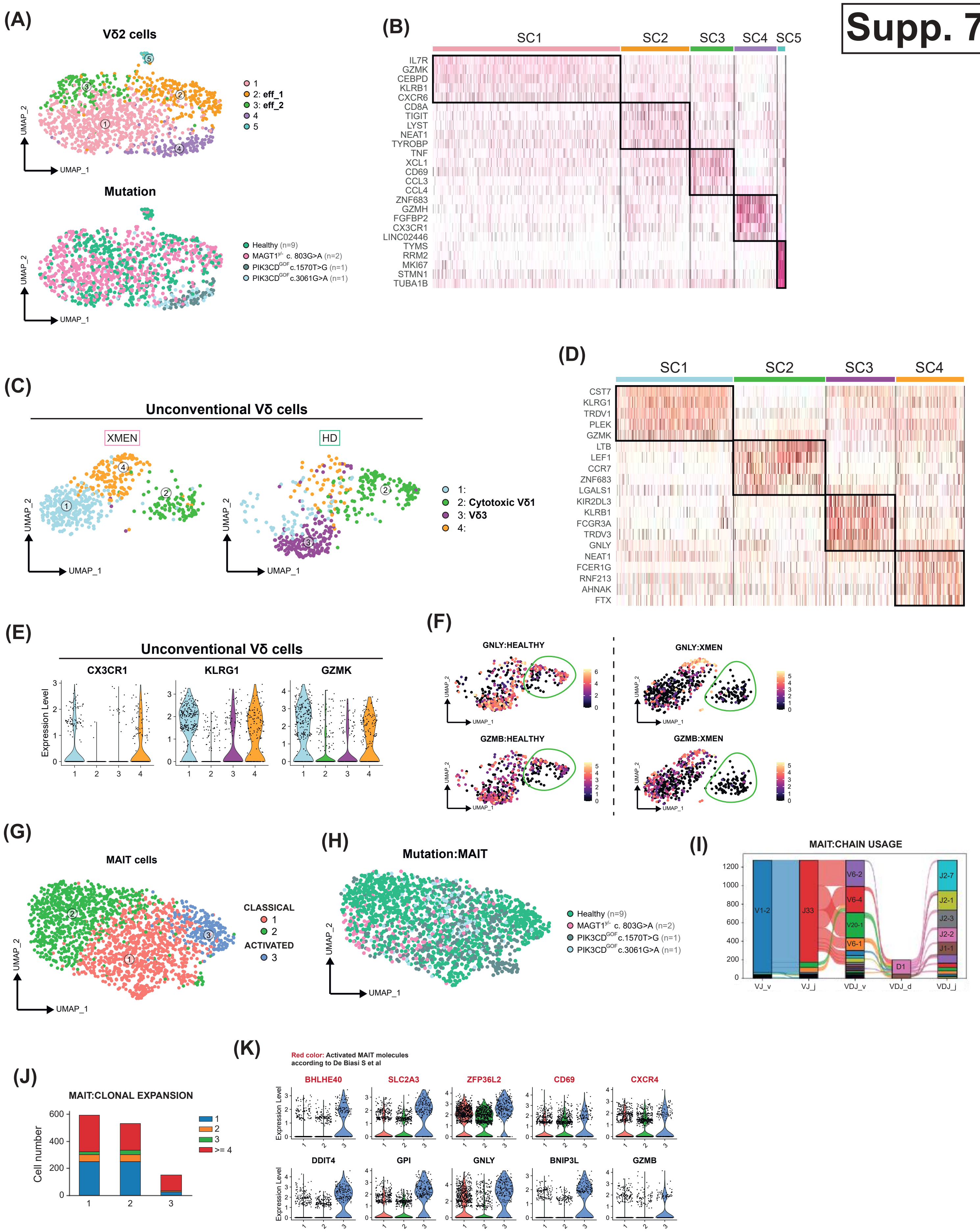
(A)
Vδ2 cells
1
2: eff_1
3: eff_2
4
5
UMAP_1
UMAP_2
Mutation
Healthy (n=9)
MAGT1y/- c. 803G>A (n=2)
PIK3CDGOF c.1570T>G (n=1)
PIK3CDGOF c.3061G>A (n=1)
(B)
SC1 SC2 SC3 SC4 SC5
IL7R
GZMK
CEBPD
KLRB1
CXCR6
CD8A
TIGIT
LYST
NEAT1
TYROBP
TNF
XCL1
CD69
CCL3
CCL4
ZNF683
GZMH
FGFBP2
CX3CR1
LINC02446
TYMS
RRM2
MKI67
STMN1
TUBA1B
(C)
Unconventional Vδ cells
XMEN
HD
1:
2: Cytotoxic Vδ1
3: Vδ3
4:
(D)
SC1 SC2 SC3 SC4
CST7
KLRG1
TRDV1
PLEK
GZMK
LTB
LEF1
CCR7
ZNF683
LGALS1
KIR2DL3
KLRB1
FCGR3A
TRDV3
GNLY
NEAT1
FCER1G
RNF213
AHNAK
FTX
(E)
Unconventional Vδ cells
CX3CR1
KLRG1
GZMK
Expression Level
(F)
GNLY:HEALTHY
GNLY:XMEN
GZMB:HEALTHY
GZMB:XMEN
(G)
MAIT cells
CLASSICAL
1
2
ACTIVATED
3
(H)
Mutation:MAIT
Healthy (n=9)
MAGT1y/- c. 803G>A (n=2)
PIK3CDGOF c.1570T>G (n=1)
PIK3CDGOF c.3061G>A (n=1)
(I)
MAIT:CHAIN USAGE
V1-2
J33
V6-2
V6-4
V20-1
V6-1
D1
J2-7
J2-1
J2-3
J2-2
J1-1
VJ_v VJ_j VDJ_v VDJ_d VDJ_j
(J)
MAIT:CLONAL EXPANSION
Cell number
1
2
3
>= 4
(K)
Red color: Activated MAIT molecules according to De Biasi S et al
BHLHE40
SLC2A3
ZFP36L2
CD69
CXCR4
DDIT4
GPI
GNLY
BNIP3L
GZMB
Expression Level

### Supp. 8

(A)

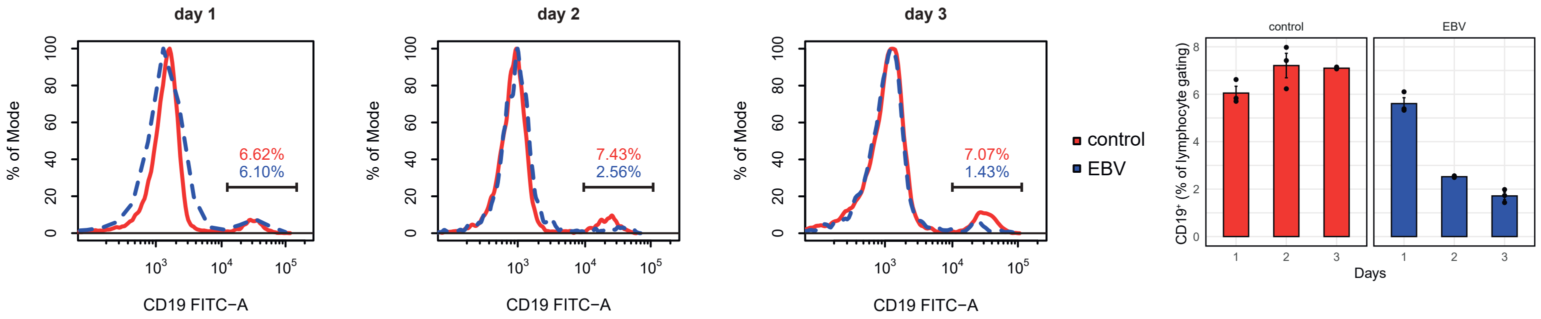


(B)

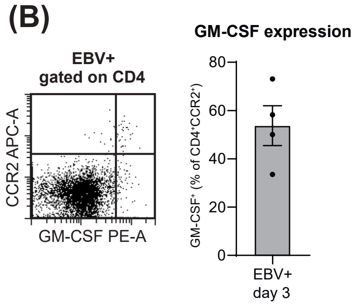


(C)

| T CELL PANEL |
|---|
| Fix Blue (UV6) |
| CD38 (NIR685) |
| CD19 (BYG667) |
| HLA-DR (APC/Cy7) |
| CD27 (FITC) |
| CD3 (PerCP5.5) |
| CD8 (PC7) |
| KLRG1 (APC) |
| CCR5 (PB) |
| CD161 (AF700) |
| CD25 (PE) |
| CD4 (v500) |
| GZMB (PE/Dazzle) |

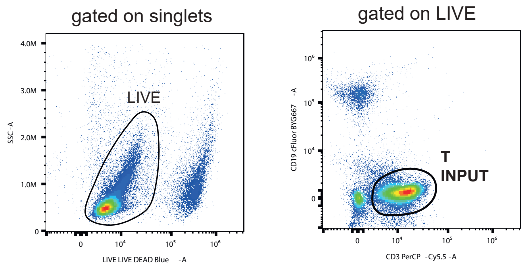


(D) (E) (F) (G)

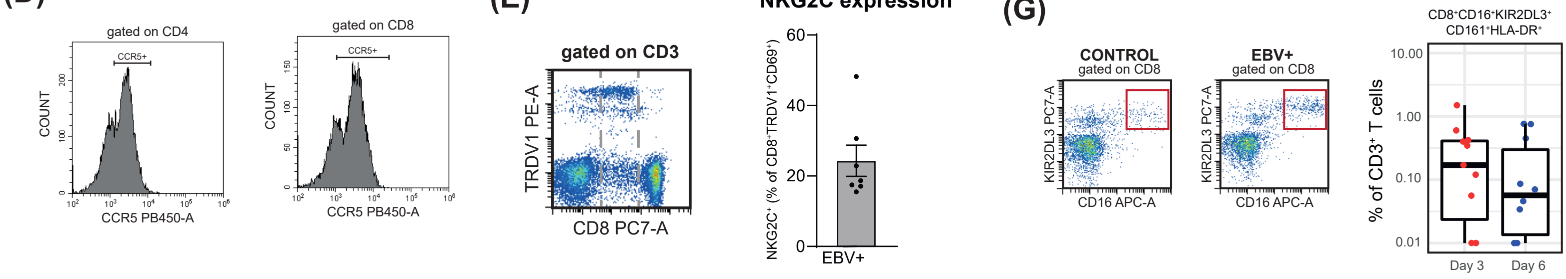


(H) (I)

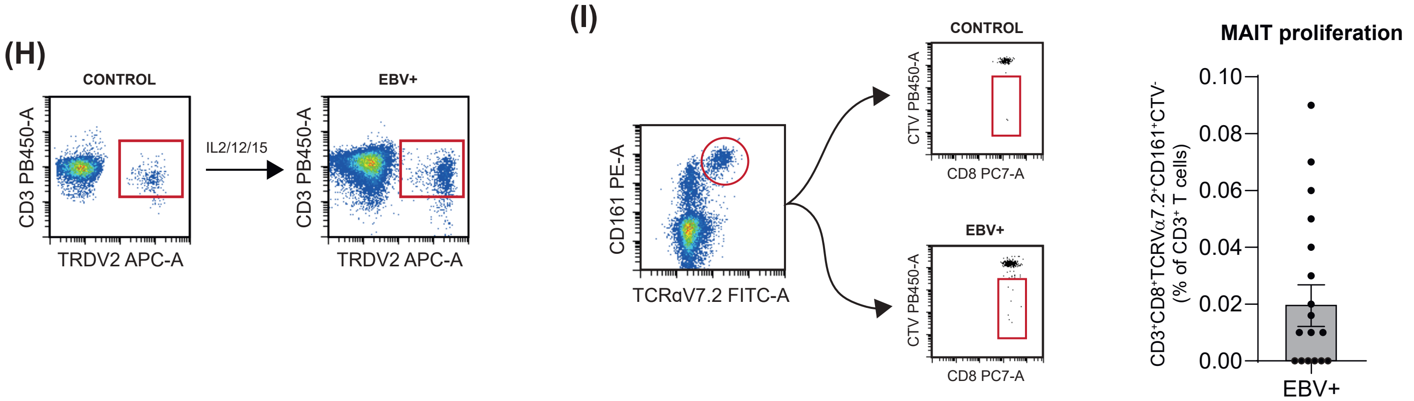


(J)

| Vδ1 \| Vδ2 \| MAIT |
|---|
| Fix Blue (UV6) |
| CD3 (PB) |
| TRDV1 (PE) |
| TRDV2 (FITC) |
| NKp30 (APC) |
| CD161 (AF700) |
| CD16 (v500) |
| HLA-DR (APC/Cy7) |
| TCRVα7.2 (PC7) |
| CD38 (NIR685) |
| GZMB (PerCP5.5) |

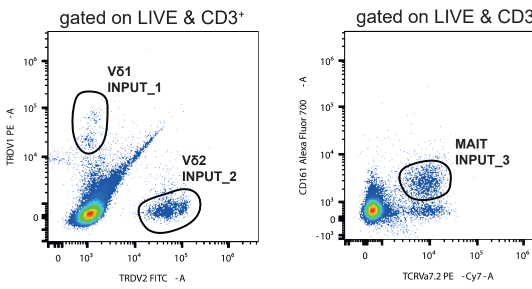


(K)

| NK PANEL |
|---|
| Fix Blue (UV6) |
| NKp80 (APC) |
| CD56 (PC5) |
| KIR (PC7) |
| CD16 (v500) |
| CD57 (PE/Dazzle) |
| HLA-DR (APC/Cy7) |
| CD94 (FITC) |
| CD3 (PB) |
| NKG2D (PE) |
| GZMB (PerCP5.5) |

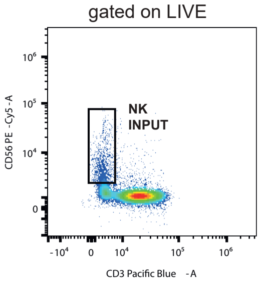

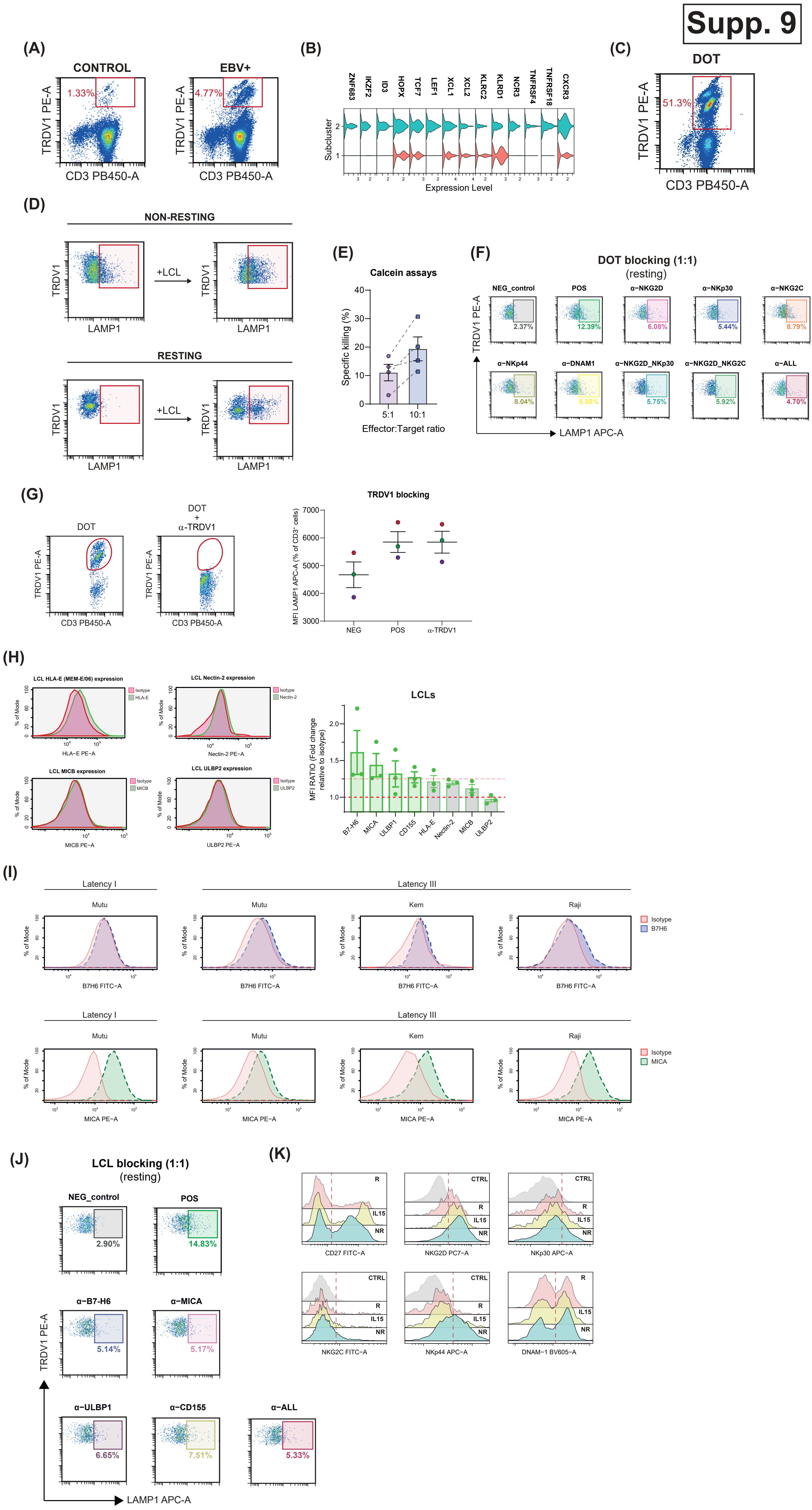

Supp. 9
(A)
CONTROL
EBV+
1.33%
4.77%
TRDV1 PE-A
CD3 PB450-A
(B)
ZNF683
IKZF2
ID3
HOPX
TCF7
LEF1
XCL1
XCL2
KLRC2
KLRD1
NCR3
TNFRSF4
TNFRSF18
CXCR3
Subcluster
Expression Level
(C)
DOT
51.3%
TRDV1 PE-A
CD3 PB450-A
(D)
NON-RESTING
RESTING
+LCL
TRDV1
LAMP1
(E)
Calcein assays
Specific killing (%)
5:1
10:1
Effector:Target ratio
(F)
DOT blocking (1:1)
(resting)
NEG_control
POS
α-NKG2D
α-NKp30
α-NKG2C
2.37%
12.39%
6.08%
5.44%
8.79%
α-NKp44
α-DNAM1
α-NKG2D_NKp30
α-NKG2D_NKG2C
α-ALL
8.04%
6.88%
5.75%
5.92%
4.70%
TRDV1 PE-A
LAMP1 APC-A
(G)
DOT
DOT + α-TRDV1
TRDV1 PE-A
CD3 PB450-A
TRDV1 blocking
MFI LAMP1 APC-A (% of CD3+ cells)
NEG
POS
α-TRDV1
(H)
LCL HLA-E (MEM-E/06) expression
LCL Nectin-2 expression
LCL MICB expression
LCL ULBP2 expression
Isotype
HLA-E
Nectin-2
MICB
ULBP2
% of Mode
HLA-E PE-A
Nectin-2 PE-A
MICB PE-A
ULBP2 PE-A
LCLs
MFI RATIO (Fold change relative to isotype)
B7-H6
MICA
ULBP1
CD155
HLA-E
Nectin-2
MICB
ULBP2
(I)
Latency I
Latency III
Mutu
Kem
Raji
Isotype
B7H6
MICA
% of Mode
B7H6 FITC-A
MICA PE-A
(J)
LCL blocking (1:1)
(resting)
NEG_control
POS
2.90%
14.83%
α-B7-H6
α-MICA
5.14%
5.17%
α-ULBP1
α-CD155
α-ALL
6.65%
7.51%
5.33%
TRDV1 PE-A
LAMP1 APC-A
(K)
CTRL
R
IL15
NR
CD27 FITC-A
NKG2D PC7-A
NKp30 APC-A
NKG2C FITC-A
NKp44 APC-A
DNAM-1 BV605-A

(A)

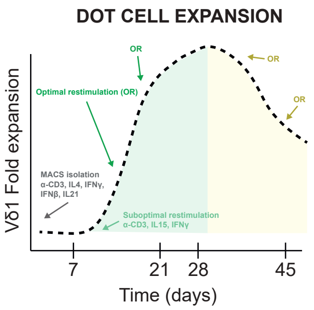


(B)

| Donor | $CD3^{+}TRDV1^{+}$ (% of $CD3^{+}$) | $CD27^{+}$ (% of $CD3^{+}TRDV1^{+}$) | $NKp30^{+}$ (% of $CD3^{+}TRDV1^{+}$) | $NKG2D^{+}$ (% of $CD3^{+}TRDV1^{+}$) | Mice injected |
|---|---|---|---|---|---|
| D1 | 21.3 | 93.5 | 31.9 | 65.4 | 2 |
| D2 | 43.3 | 58.8 | 26.1 | 59.3 | 1 |

(C)

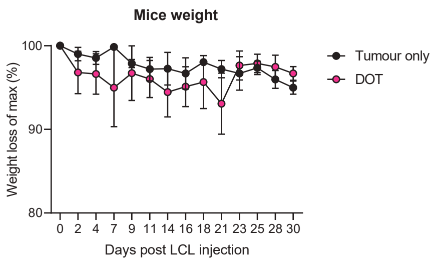


(D)

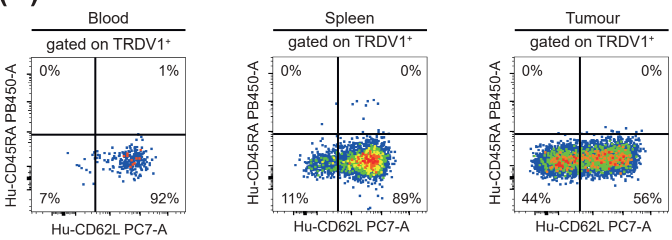


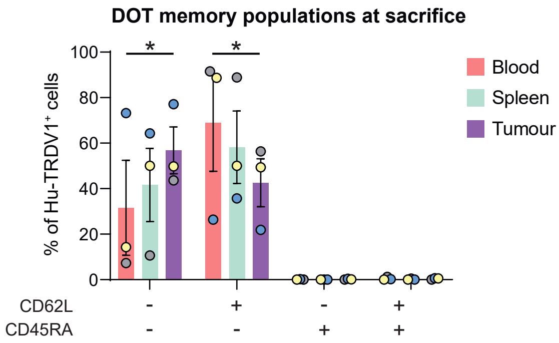


(E)

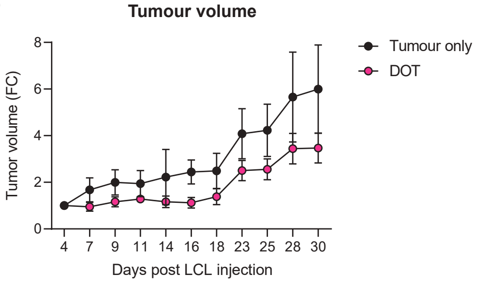

### Supp. 11

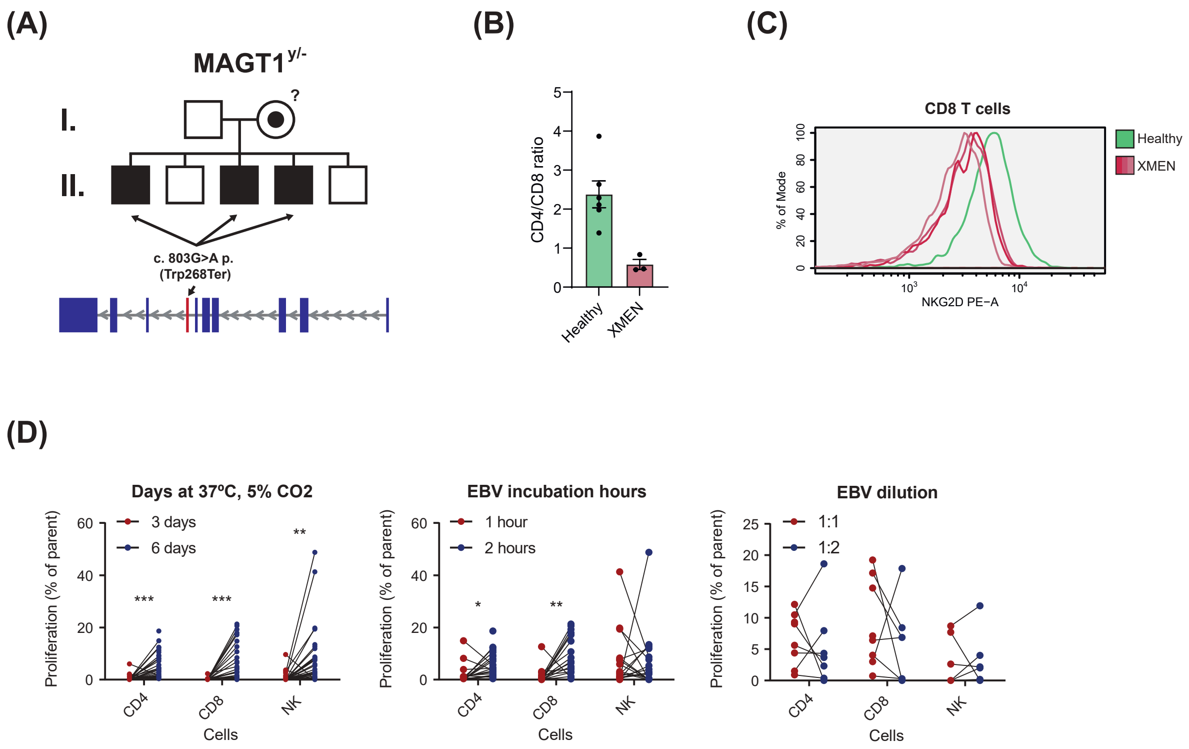
(A)
MAGT1 y/-
I.
?
II.
c. 803G>A p. (Trp268Ter)
(B)
CD4/CD8 ratio
Healthy
XMEN
(C)
CD8 T cells
% of Mode
NKG2D PE-A
Healthy
XMEN
(D)
Days at 37ºC, 5% CO2
3 days
6 days
***
***
**
EBV incubation hours
1 hour
2 hours
*
**
EBV dilution
1:1
1:2
Proliferation (% of parent)
CD4
CD8
NK
Cells
